# Ubiquitination steers SRF3 plasma membrane nano-organization to specify signaling outputs

**DOI:** 10.1101/2022.10.28.514292

**Authors:** Platre Matthieu Pierre, Gleason Flores Matias, Brent Lukas, Cao Min, Zhang Ling, Santosh B. Satbhai, Neveu Julie, Vert Gregory, Busch Wolfgang

## Abstract

Organisms cope with myriads of competing and conflicting environmental signals. These signals are often perceived by cell surface receptor kinases to mount appropriate adaptive responses. However, it is not well understood by which mechanism single receptor kinases can transduce different signals. The plant receptor kinase SRF3 transduces low iron and bacteria-derived signals. We found that upon these signals, ubiquitinated SRF3 is recognized by clathrin-mediated endocytosis for vacuolar targeting. Live super resolution microscopy revealed that cell surface SRF3 is present in a fast diffusible fraction, which is sustained by ubiquitination, and that non-ubiquitinated SRF3 is present in immobile nanodomains. Ubiquitination-mediated degradation of SRF3 is required for signaling only under low iron but not upon flg22 perception. Flg22-triggered SRF3 phosphorylation leads to SRF3 accumulation in the immobile fraction in which degradation is restricted, thereby preventing low iron signaling. We therefore propose that ubiquitination-dependent plasma membrane nano-organization of SRF3 specifies its signal transduction pathways.

## INTRODUCTION

Plants are sessile organisms which need to constantly adapt to their everchanging environment to thrive and survive. To adapt their physiology and development, they have evolved sophisticated sensing and signaling systems to perceive and integrate external environmental signals. Key components of these systems are cell surface receptor kinases that transduce external signals and elicit intracellular signaling via effectors proteins to fine tune cellular responses^1^. For example, the plant leucine-rich repeat receptor kinase (LRR-RK) FLAGELLIN SENSING 2 (FLS2) is able to recognize bacterial presence by binding the last 22 amino acids of the bacterial flagellin protein (flg22)^2^. This recognition is followed by a direct interaction with the co-receptor BRASSINOSTEROID INSENSITIVE 1-ASSOCIATED KINASE 1/SOMATIC EMBRYOGENESIS RECEPTOR KINASE 3 (BAK1/SERK3) to trigger a phosphorylation cascade that is critical for plasma membrane defense signaling through the intracellular effector protein, BIK1^3,4^. In some cases, co-receptors are involved in transducing different signaling pathways depending on the signal perceived. For instance, BAK1/SERK3 is also required to trigger phosphorylation-dependent signaling by interacting directly with BRASSINOSTEROID INSENSITIVE 1 (BRI1) in the presence of the plant hormone brassinosteroids to activate growth through the intracellular effector proteins BSK1/CDG1^5,6^. Thus, the BAK1/SERK3 co-receptor has been proposed to be a key stone in balancing growth-defense responses in plants^7^. However, how a co-receptor can interact specifically with different receptors to promote different signaling pathways remain still unclear. Based on singe particle tracking, it has been proposed that BRI1 and FLS2 segregate in distinct nanodomains at the plasma membrane, thereby allowing signaling specificity while relying on the same co-receptor^8^. Additionally, a recent study demonstrated that specific phosphosites on BAK1 are phosphorylated only under flg22 elicitation but not under brassinosteroids perception which in turn specify signaling^9^. Another example for two pathways being targeted by the same membrane protein is the small GTPase Rho of plant 6 (ROP6) that signals under osmotic stress and in the presence of the phytohormone auxin. It is located within specific plasma membrane nanodomains, which are key to specify signaling^10,11^. While the receptor kinase involved in auxin perception and signal transduction through ROP6 has been discovered, the link between ROP6 and the upstream osmotic receptors OSCA1 still remain unclear^12,13^. Functionally dichotomous roles of receptors have been also observed in animals. For instance, the plasma membrane-associated EPIDERMAL GROWTH FACTOR RECEPTOR (EGFR) receptor kinase is situated at the nexus of two pathways. These AKT and MEK pathways are involved in the regulation of cell growth and survival, two outputs determining oncogenic cell formation^14,15^. In this case, the small GTPase K-ras, which is downstream of EGFR, is depending on its polybasic domain and prenyl anchor, to be addressed into specific lipid-composition nanodomains at the plasma membrane to balance signaling^16^. While it is therefore clear that the allocation of signaling proteins to dedicated nanodomains specifies signaling in several eukaryotic systems, it is not well understood how receptor kinases are localized to distinct regions of the plasma membrane.

Many receptors (e.g. BRI1 and FLS2) and co-receptors (e.g. BAK1) are endocytosed to fine tune phosphorylation-dependent signaling upon signal perception. This removal from the plasma membrane is mainly mediated by the clathrin-dependent machinery ^17–20^. Regulation of receptor kinases endocytosis must be tightly regulated. One way by how endocytosis rates are determined is via post translational protein modification. For example, EGFR, BRI1 and FLS2 are ubiquitinated ^17,21,22^. BRI1 and EGFR signaling is tuned by the addition of K63 ubiquitin chains which trigger its removal from the plasma membrane and subsequent vacuolar/lysosomal targeting through endosomal trafficking^17,21^. Moreover, the putative plant glucose D-receptor AtRGS1 is endocytosed by either the clathrin-dependent or independent machinery to specify the signaling pathway of the conserved G protein in presence of glucose and the bacterial elicitor flg22^23^. Overall, to specify and fine tune signaling receptor/co-receptor complexes can induce differential phosphorylation, ubiquitination, endocytosis and be localized into specific nanodomains at the plasma membrane. Nonetheless, how these factors interplay to specify signaling remains unexplored.

Iron homeostasis and defense pathways are tightly intertwined in eukaryotes given the micronutrient iron is critical for determining host-pathogen interactions^24^. Iron has been proposed to be involved in mediating host nutritional immunity and as a priming signal to initiate host defense responses^25–29^. The STRUBBELIG-RECEPTOR FAMILY 3 (SRF3) LRR-RK has been recently shown to mediate low iron and flg22-dependent root growth and transcriptional signaling, placing this receptor at the nexus of both pathways^27^. An important part of the SRF3 signaling mechanism is its decreased abundance at the plasma membrane in response to low iron and flg22-derived signals^27^. Here, we present the molecular mechanism by which SRF3 can trigger specific signaling pathways and how it relates to it abundance and localization at the plasma membrane. We found that ubiquitination-triggered SRF3 clathrin-dependent endocytosis is key for this regulation which leads to its vacuolar degradation upon both signals. Using super resolution microscopy, we showed that cell surface SRF3 segregates into mobile and immobile fractions, which are dependent on SRF3 ubiquitination status. Ubiquitinated SRF3 signals from the mobile fraction are required to regulate low iron responses while non-ubiquitinated SRF3 signals from the immobile nanodomains are required to regulate flg22 responses. Accordingly, a SRF3 flg22-phosphominic version was localized in the immobile fraction and prevented its degradation. Our work highlights the capacity of post-translational modifications such as ubiquitination to specify the signaling output of receptor kinases by affecting their nanoscale organization.

## RESULTS

### SRF3 accumulates along the endocytic pathway and is targeted to the vacuole under low iron levels and flg22

Recently, we had found that *SRF3* is critical in regulating the root growth response to low iron levels and to the bacterial peptide flagellin 22 (flg22)^27^. Under both conditions SRF3 is rapidly degraded from the plasma membrane (Supplementary Data 1a)^27^. We set out to explore by which cellular mechanism SRF3 is degraded from this compartment. When looking at fluorescently tagged SRF3 in unstressed condition, we noticed that it accumulates in intracellular dots and large intracellular structures, suggesting its presence in endosomal compartments and vacuole, thereby resembling the endocytic pathway (Supplementary data 1b). To test this assumption, we assessed the accumulation of SRF3 at different steps of the endocytosis pathway from the plasma membrane to the vacuole. Consistent with our hypothesis, SRF3 co-localized with the plasma membrane marker Lti6b (Supplementary data 1c). Moreover, the early endosomes trans-golgi network (EE/TGN) marker, VTI12, revealed a partial colocalization with SRF3 (Supplementary Data 1d). To allow an easy visualization of this compartment, we applied Brefeldin A (BFA) which inhibits protein recycling and to a lesser extent endocytosis to gather the plasma membrane-derived early endosomes trans-golgi network (EE/TGN) into the so-called BFA body^30–32^. Under this treatment, we observed a co-localization of SRF3 and the EE/TGN marker VTI12 in BFA bodies further confirming that SRF3 is present in this compartment (Supplementary Data 1e). Colocalization analysis of SRF3 with the late endosomal (LE) marker Rab F2, revealed also a partial colocalization (Supplementary Data 1f). To corroborate this finding, we applied wortmannin (Wm) to facilitate LE visualization as this treatment leads to easily recognizable donut shape-like late endosomes structures^30,32^. In this condition, Rab F2 also colocalized partially with SRF3 showing that a fraction of SRF3 protein accumulates within this compartment (Supplementary Data 1g). Finally, we assessed the presence of SRF3 in lytic compartments by using plants coexpressing the tonoplast marker VHA-A3 with SRF3 and placing the plants in the dark in order to increase GFP stability in the vacuole^33^. This revealed that SRF3 accumulated also in the vacuole (Supplementary Data 1h). Taken together, SRF3 accumulates along the endocytic pathway under unstressed conditions.

As low iron conditions and treatment with flg22 cause a rapid removal of SRF3 from the plasma membrane^27^, we wanted to test whether this decrease could be caused by an increase of SRF3 endocytosis-triggered vacuolar targeting. We therefore evaluated the size of the BFA body as a proxy to evaluate the enrichment of SRF3 in EE/TGN^10^. We applied a concomitant treatment of BFA with low iron (−Fe) or low iron supplemented with the iron chelator Ferrozine at 100 μM (−FeFZ[100]) or with 1 μM of flg22[1] or flg20[1] for which the two last amino acids of flg22 were removed to inactivate the peptide. The size of BFA bodies under both low iron conditions and flg22 treatment were larger compared to their controls and showed an enrichment of SRF3 within EE/TGN compartments (Fig. 1a,d). To further corroborate that both signals trigger SRF3 enrichment along the endocytic pathway, we applied Wm and counted the number of donut shape-like late endosomes. Under both treatments more Wm-positive late endosomes were counted compared to the controls (Fig. 1b,e). This supported the model that SRF3 is trafficked to EE/TGN and LE at increased rates in response to low iron or flg22, thereby displaying a faster endocytosis rate en route to the vacuole (Fig. 1b,e). To further test this model, we determined SRF3 levels within the vacuole by placing plants in the dark during the application of low iron levels or flg22. More intracellular signal was detected compared to the control in both conditions, showing that SRF3 is further enriched in the vacuole under these conditions (Fig. 1c,f). Overall, these experiments support that SRF3 is constitutively endocytosed and that low iron levels or flg22 further stimulate SRF3 endocytosis to mediate an increased targeting in the vacuole therefore removing SRF3 from the plasma membrane.

**Fig. 1:**
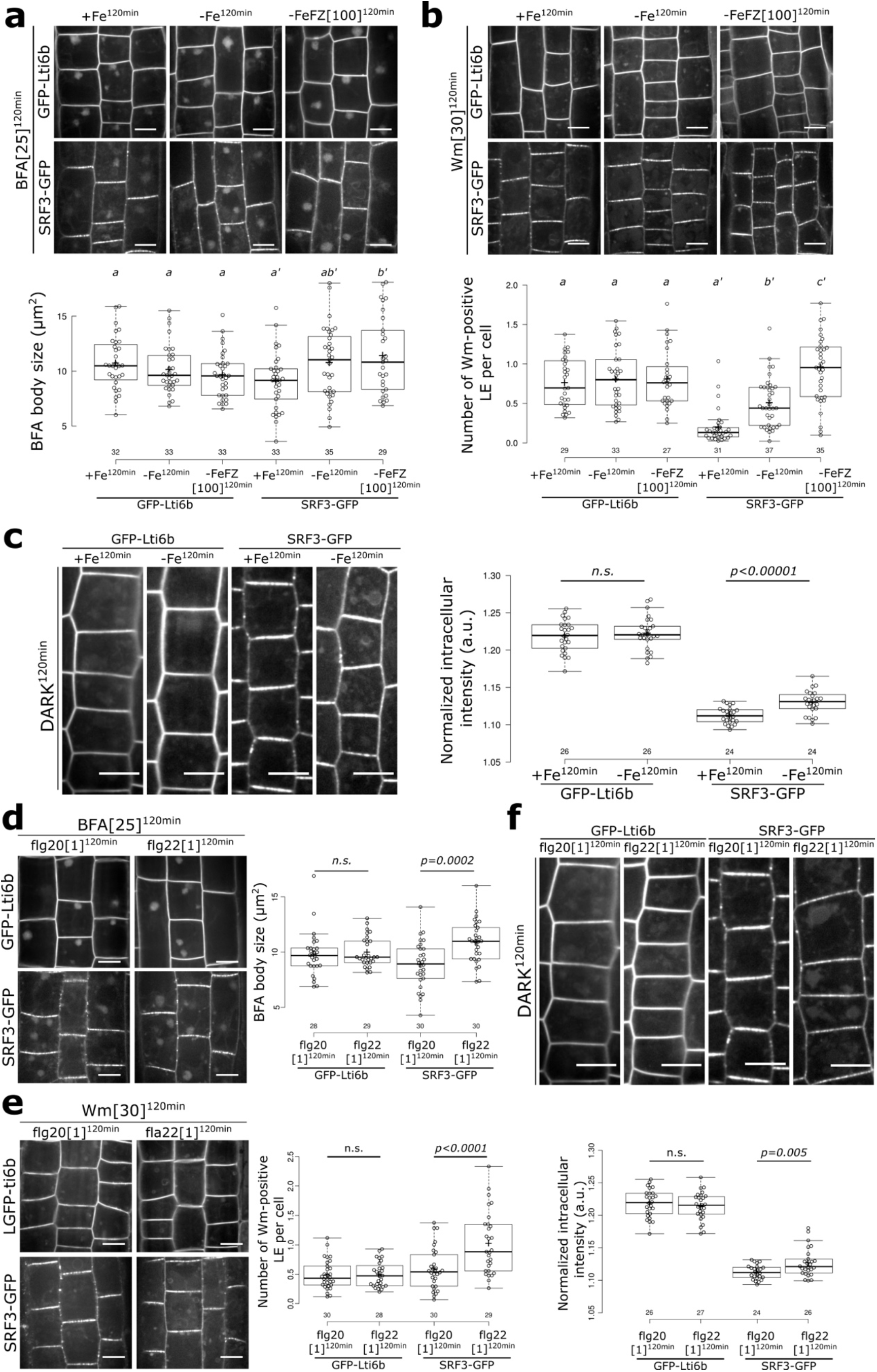
Endosomal trafficking targets SRF3 to the vacuole under low iron and flg22. **a**, Upper panel: confocal images of root epidermal cells of 5-day old seedlings expressing *pSRF3::SRF3-GFP* (SRF3-GFP) *or p35s::GFP-Lti6b* (GFP-Lti6b) co-treated with BFA at 25μM and +Fe, −Fe and −Fe supplemented with Ferrozine (−FeFZ) at 100 μM for 120 minutes. Lower panel: quantification of the BFA body size (μm). Scale bar, 5μm. [two-way ANOVA followed by a post hoc Tukey HSD test; letters indicate statistical differences (p<0.05)]. **b**, Upper panel: confocal images of root epidermal cells of 5-day old seedlings expressing *pSRF3::SRF3-GFP* (SRF3-GFP) *or p35s::GFP-Lti6b* (GFP-Lti6b) co-treated with Wm at 30 μM and +Fe, −Fe and −Fe supplemented with Ferrozine (−FeFZ) at 100 μM for 120 minutes. Lower panel: quantification of the number of Wm-positive endosomes (a.u). Scale bar, 5μm. [two-way ANOVA followed by a post hoc Tukey HSD test; letters indicate statistical differences (p<0.05)]. **c**, Left panel: confocal images of root epidermal cells of 5-day old seedlings expressing *pSRF3::SRF3-GFP* (SRF3-GFP) *or p35s::GFP-Lti6b* (GFP-Lti6b) treated +Fe, −Fe and −Fe supplemented with Ferrozine (−FeFZ) at 100 μM and placed in the dark for 120 minutes. Right panel: quantification of the normalized intracellular intensity (a.u.). Scale bar, 5μm. [Independent two-way Student’s T-test (p<0.05)]. **d**, Left panel: confocal images of root epidermal cells of 5-day old seedlings expressing *pSRF3::SRF3-GFP* (SRF3-GFP) *or p35s::GFP-Lti6b* (GFP-Lti6b) co-treated with BFA at 25μM and flg20 or flg22 at 1 μM for 120 minutes. Right panel: quantification of the BFA body size (μm). Scale bar, 5μm. [Independent two-way Student’s T-test (p<0.05)]. **e**, Left panel: confocal images of root epidermal cells of 5-day old seedlings expressing *pSRF3::SRF3-GFP* (SRF3-GFP) *or p35s::GFP-Lti6b* (GFP-Lti6b) cotreated with Wm at 30 μM and flg20 or flg22 at 1 μM for 120 minutes. Right panel: quantification of the number of Wm-positive endosomes (a.u.). Scale bar, 5μm. [Independent two-way Student’s T-test (p<0.05)]. **f**, Upper panel: Confocal images of root epidermal cells of 5-day old seedlings expressing *pSRF3::SRF3-GFP* (SRF3-GFP) *or p35s::GFP-Lti6b* (GFP-Lti6b) treated with flg20 or flg22 at 1 μM and placed in the dark for 120 minutes. Right panel: quantification of the normalized intracellular intensity (a.u.). Scale bar, 5μm.

### Clathrin-dependent endocytosis controls SRF3 plasma membrane levels in response to low iron and flg22

We then wondered which type of endocytosis is involved in SRF3 removal from the plasma membrane. The main route mediating cargo protein endocytosis is through the clathrin machinery. To be able to observe which endocytosis mechanism is relevant for SRF3, we utilized variable angle epifluorescence microscopy based on total internal reflection fluorescence microscopy (VAEM-TIRF), which allows observations in close vicinity of the plasma membrane (~100 nm)^34^. This microscopy technique revealed a partial colocalization between clathrin light chain 2 (CLC2) with SRF3 at the plasma membrane, suggesting an involvement of clathrin in SRF3 endocytosis (Fig. 2a). To test this hypothesis, we took a genetic approach using an estradiol inducible AUXILIN-LIKE 2 version (*XVE>>AX2*) to inhibit clathrin-dependent endocytosis while monitoring SRF3 plasma membrane levels^35^. After 18 hours of induction, we detected an increase of the plasma membrane mean gray value (Fig. 2b), highlighting that SRF3 clathrin-dependent endocytosis modulates SRF3 plasma membrane levels. To test whether clathrin-dependent endocytosis is critical in regulating SRF3 plasma membrane degradation upon low iron levels and flg22, we utilized the clathrin heavy chain 1 and 2 mutants, *chc1-1* and *chc2-3*^36^. While in control conditions, we did not observe any differences in SRF3 accumulation at the plasma membrane, under low iron levels and upon flg22 treatment only a slight decrease was noticed but to a lesser extent compared to the WT (Fig. 2c-d). Moreover, to evaluate SRF3 intracellular trafficking in endosomes and vacuole, we calculated SRF3 normalized intracellular intensity which corresponds to SRF3 cytosolic signal intensity divided by SRF3 total signal intensity. This parameter was not drastically changed in basal conditions. (Supplementary Data 2a-b). Furthermore, in contrast to WT, we did not observe an increase of the normalized intracellular intensity in *chc* mutants under low iron levels and flg22, supporting our hypothesis that clathrin is required to mediate SRF3 intracellular trafficking and enrichment in the vacuole under low iron of flg22 treatments (Supplementary Data 2a-b). To further test the involvement of clathrin in triggering SRF3 endocytosis, we applied endosidin9-17 (ES9-17) a well characterized clathrin heavy chain inhibitor^37^ upon low iron and flg22 treatments. Consistent with our results from the *chc* mutants, neither low iron nor flg22 were able to decrease SRF3 plasma membrane levels upon ES9-17 compared to mock condition (Fig. 2e). Altogether, our data demonstrate that SRF3 is to a lesser extent endocytosed by clathrin-mediated endocytosis in standard conditions, but it is critical when plants are exposed to decreased iron levels or flagellin peptide.

**Fig. 2:**
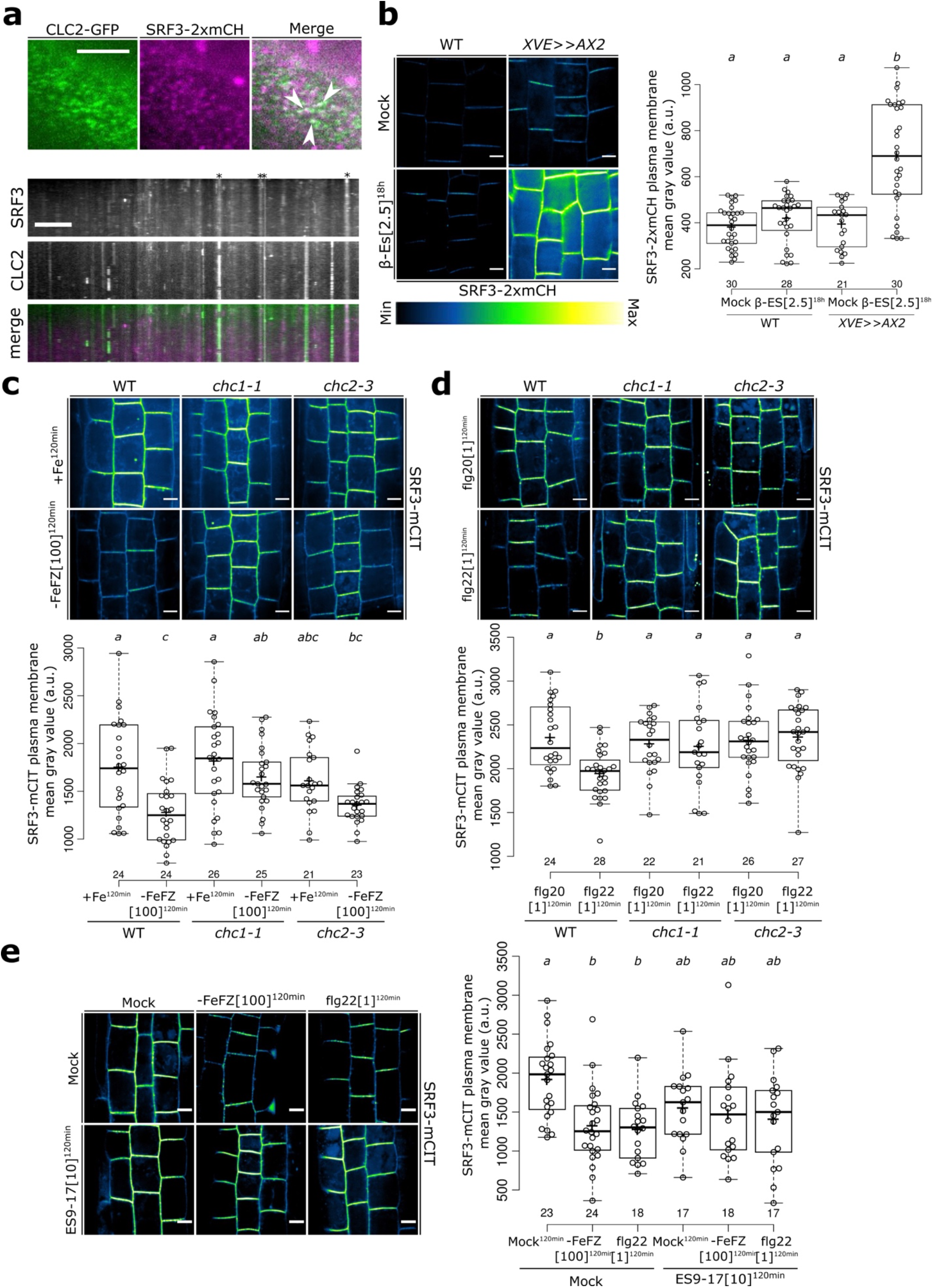
Clathrin-dependent endocytosis regulates SRF3 levels at the plasma membrane upon low iron and flg22. **a**, Upper panel: micrographs of 5-day old seedlings of root epidermal cells co-expressing *pCLC2::CLC2-GFP* (CLC2-GFP) and *pUBQ10::SRF3-2xmCHERRY* (SRF3-2xmCH). Lower panel: representative kymograph. Scale bar, 1μm. **b**, Left panel: confocal images of root epidermal cells of 5-day old seedlings expressing *pUBQ10::SRF3-2xmCHERRY* or co-expressing *pUBQ10::SRF3-2xmCHERRY* (SRF3-2xmCH) and *pUBQ10-XVE::AUXILIN-LIKE1 (XVE>>AX2)* under mock treatment or under β-Estradiol at 2.5μm (β-ES[2.5]) for 18 hours. Right panel: quantification of SRF3 plasma membrane mean gray value (a.u.). Scale bar, 5μm. [two-way ANOVA followed by a post hoc Tukey HSD test; letters indicate statistical differences (p<0.05)]. **c**, Upper panel: confocal images of root epidermal cells of 5-day old seedlings expressing *pUBQ10::SRF3-mCITRINE* (SRF3-mCIT) in WT, *chc1-1* or *chc2-3* mutants treated with +Fe or −Fe supplemented with Ferrozine (−FeFZ) at 100 μM for 120 minutes. Lower panel: quantification of SRF3 plasma membrane mean gray value (a.u.). Scale bar, 5μm. [two-way ANOVA followed by a post hoc Tukey HSD test; letters indicate statistical differences (p<0.05)]. **d**, Upper panel: confocal images of root epidermal cells of 5-day old seedlings expressing *pUBQ10::SRF3-mCITRINE* (SRF3-mCIT) in WT, *chc1-1* or *chc2-3* mutants treated with flg20 or flg22 at 1 μM for 120 minutes. Lower panel: quantification of SRF3 plasma membrane mean gray value (a.u.). Scale bar, 5μm. [two-way ANOVA followed by a post hoc Tukey HSD test; letters indicate statistical differences (p<0.05)]. **e**. Left panel: confocal images of root epidermal cells of 5-day old seedlings expressing *pUBQ10::SRF3-mCITRINE* (SRF3-mCIT) co-treated with ES9-17, mock, −Fe supplemented with Ferrozine (−FeFZ) at 100 μM or flg22 at 1μM for 120 minutes. Right panel: quantification of SRF3 plasma membrane mean gray value (a.u.). Scale bar, 5μm. [two-way ANOVA followed by a post hoc Tukey HSD test; letters indicate statistical differences (p<0.05)].

### Ubiquitination regulates SRF3 plasma membrane levels and its targeting to the vacuole in response to low iron and flg22

To further dissect the role of SRF3 endocytosis, we set out to identify which molecular mechanism triggers SRF3 endocytosis. SRF3 is member of the Strubbelig Receptor kinase family, among them SRF6, 7, 8 and 9 that have been shown to be ubiquitinated and it was predicted that it occurred at residues localized in the juxtamembrane domain (JD)^38,39^. Using four different prediction websites for protein ubiquitination, we identified 5 lysine residues as possible ubiquitination sites in the SRF3 JD that were predicted with high-confidence or by two or more of these algorithms (Fig. 3a). Mutation of the predicted ubiquitinated residues into arginine (R) stabilized SRF3 (SRF3^5KR^) levels in cells compared to the wild type version of SRF3 (SRF3^WT^) as observed by western blot using anti-GFP antibody when compared to Tubulin loading (Supplementary Data 3b). Importantly, the mRNA levels were indistinguishable in these lines (Supplementary Data 3a-b), demonstrating that the effect on protein levels were due to the lysine ubiquitination. However, quantification of the mean gray values in the root tip (meristematic and elongation zones, excluding the root cap) by confocal microscopy did not reveal an increase of protein abundance in this zone (Supplementary Data 3c). We found that this increase was due to an SRF3 over-accumulation in the differentiation zone as indicated by the ratio of the mean gray value of the root tip with the differentiation zone (Supplementary Data 3d). Co-immunoprecipitation using an antibody against general ubiquitination revealed that SRF3^5KR^ was less ubiquitylated compared to the WT form (Fig. 3b and Supplementary Data 3e). These results show that SRF3 is constitutively ubiquitinated, which tunes its degradation rate. We then wondered whether SRF3 ubiquitination is required to regulate SRF3 protein levels and degradation under low iron levels and flg22 in the root transition/elongation zone. Confocal microscopy experiments revealed that SRF3^WT^ but not SRF3^5KR^ is degraded at the plasma membrane under these conditions compared to control conditions (Fig. 3c). Coimmunoprecipitation (Co-IP) using an antibody against general ubiquitination seemed to indicate a slight increase of SRF3 ubiquitination in low iron and flg22 (Fig. 3b). Taken together these results show that SRF3 ubiquitination regulates its plasma membrane levels under flg22 and low iron levels.

**Fig. 3:**
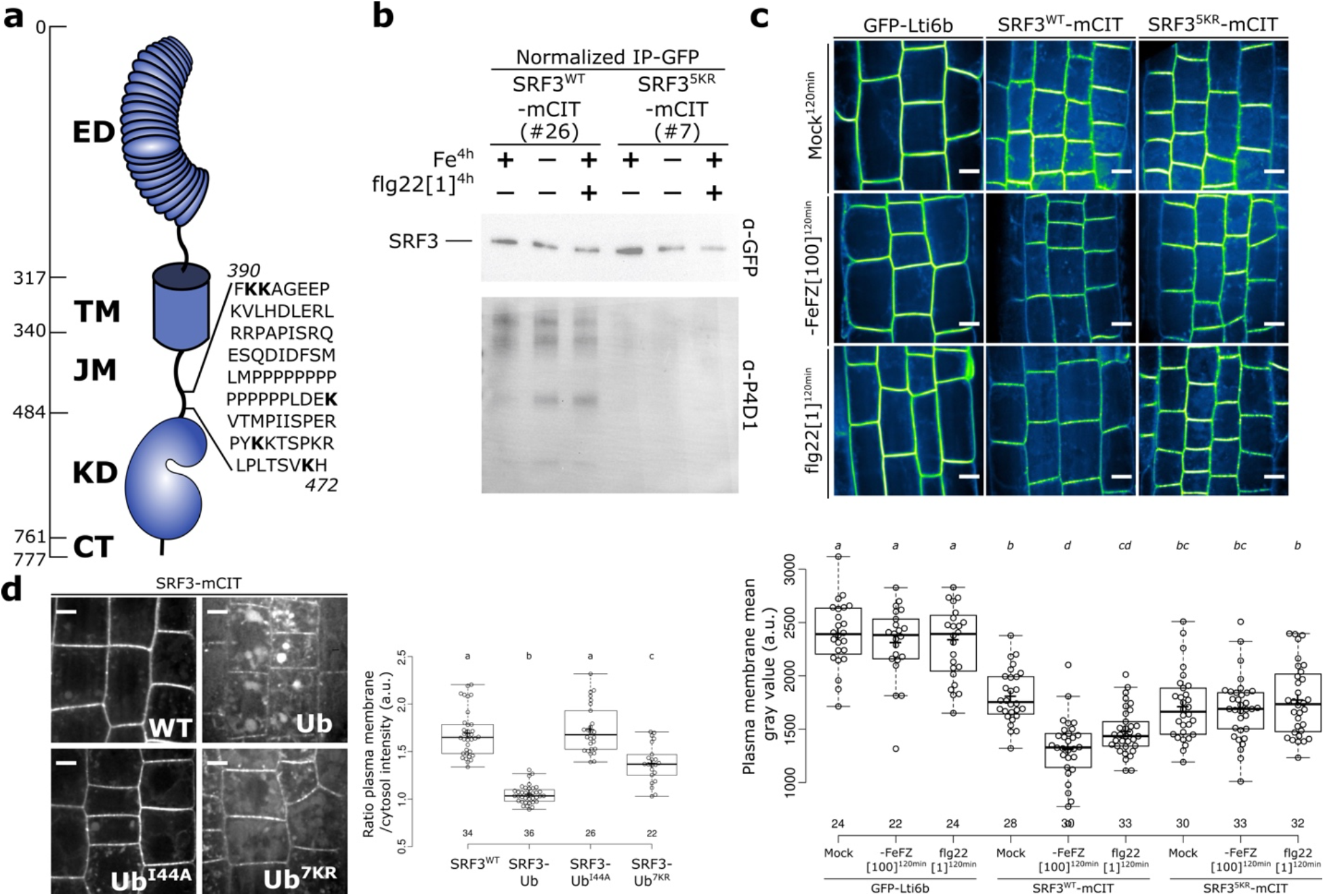
Ubiquitination in the juxta-membrane domain of SRF3 triggers degradation upon low iron and flg22. **a**, Schematic representation of SRF3. **b**, Co-immunoprecipitation (Co-IP) using α-GFP on root material from 12-day old seedlings expressing *pUBQ10::SRF3^WT^-mCITRINE* (SRF3^WT^-mCIT) or *pUBQ10::SRF3^5KR^-mCITRINE* (SRF3^5KR^-mCIT) in standard conditions or treated with −Fe and flg22 at 1μM for 4 hours where the input has been normalized to obtain the same protein quantity and on which the ubiquitination is determined using α-P4D1. **c**, Upper panel: confocal images of root epidermal cells of 5-day old seedlings expressing *pUBQ10::SRF3^WT^-mCITRINE* (SRF3^WT^-mCIT) or *pUBQ10::SRF3^5KR^-mCITRINE* (SRF3^5KR^-mCIT) or *p35s::GFP-Lti6b* (GFP-Lti6b) under mock, −Fe supplemented with Ferrozine (−FeFZ) at 100 μM or flg22 at 1μM for 120 minutes. Lower panel: quantification of SRF3 plasma membrane mean gray value (a.u.). [two-way Kruskal-Wallis coupled with post hoc Steel-Dwass-Critchlow-Fligner procedure (p<0.05)]. **d**, Left panel: confocal images of root epidermal cells of 5-day old seedlings expressing *pUBQ10::SRF3^WT^-mCITRINE* (*SRF3^WT^*), *pUBQ10::SRF3-mCITRINE^Ub^* (*SRF3^Ub^*), *pUBQ10::SRF3-mCITRINE^Ub-144A^* (*SRF3^Ub-144Ab^*), and pUBQ10::SRF3-mCITRINE^Ub-7KR^ (*SRF3^Ub-7KR^*). Right panel: quantification of the ratio of plasma membrane/cytosol intensity (a.u.). Scale bar, 5μm. [two-way ANOVA followed by a post hoc Tukey HSD test, letters indicate statistical differences (p<0.05)].

We had shown that low iron levels and flg22 trigger SRF3 trafficking to reach the vacuole (Figure 1). Since poly-ubiquitination is involved in protein targeting to the vacuole, we tested if SRF3 can be targeted to these lytic compartments by this mechanism^40^. For that, we generated three lines in which SRF3 was artificially ubiquitylated after the fluorescent tag and determined SRF3 cellular enrichment by calculating the plasma membrane/cytosol mean gray value ratio of SRF3^Ub^, SRF3^Ub-144A^ and SRF3^Ub-7KR^ compared to SRF3^WT^ (Supplementary Data 3f). SRF3^Ub^ for which a ubiquitin moiety was added, showed a decrease of the ratio indicating low SRF3 levels at the plasma membrane and at the same time showed a pronounced accumulation in the vacuole and intracellular compartments compared to SRF3^WT^ (Fig. 3d). To further test whether SRF3 targeting to the vacuole was indeed triggered by the addition of the Ub moiety, we analyzed the ratio of a plant transformed with a SRF3^Ub-144A^ version, which prevents the recognition of the ubiquitin moiety by the ubiquitin machinery^40^. This version presented a similar plasma membrane/cytosol ratio value as SRF3^WT^ (Fig. 3d). Finally, for a plant line containing SRF3^Ub-7KR^, which prevents poly-ubiquitination^40^, we noticed a slight decrease of the ratio compared to SRF3^WT^ but it was not as pronounced as for SRF3^Ub^ (Fig. 3d). This corroborated that SRF3 poly-ubiquitination targets SRF3 from the plasma membrane to the vacuole. However, it is possible that another type of ubiquitination might occur in this version, as the ratio observed for SRF3^Ub-7KR^ was still lower compared to that of SRF3^WT^. Taken together our data suggest that upon low iron and flg22 SRF3 is polyubiquitinated, which then targets SRF3 to the vacuole and therefore mediates its degradation.

### SRF3 is organized into immobile nanodomains and a diffusible fraction at the plasma membrane

We had found that SRF3 proteins co-exist at the plasma membrane in two different states prior to be endocytosed, an un-ubiquitinated and a ubiquitinated form. We therefore wanted to investigate the role of SRF3 ubiquitination on its plasma membrane dynamics and organization. For this, we applied the super resolution microscopy modality single particle tracking photoactivated localization microscopy (Spt-PALM)^41^. To do this, we generated a version of SRF3 that was fused to the photoconvertible protein mEOS2 and driven by the *UBIQUITIN10* promotor (SRF3-mEO2S). In standard conditions, we observed the presence of two populations describing a normal distribution (Supplementary Data 4a). We separated different fractions using a mixture model of scale mixtures of skew-normal denoted as FMSMSN. Using this approach, we were able to separate two different fractions by a log(D) of −1.67 (Supplementary Data 4a)^41^. As non-mobile particles should have a low log(D) value, we reasoned that the fraction above −log(D) of 1.67 corresponded to a mobile fraction of SRF3, while the fraction below this represented an immobile fraction. 70% of SRF3^WT^ is present in the immobile fraction and about 30% in the mobile fraction (Fig. 4a-b). We then set out to perform clustering analysis on the immobile fraction using a Tessellation approach to determine whether this fraction is organized randomly or aggregates in some specific domains at the plasma membrane^42^. We found that about 10% of SRF3^WT^ immobile fraction accumulates in nanodomains displaying a diameter of ~180nm (Fig. 4d-e and Supplementary Data 4b). We therefore conclude that under standard conditions, a majority of SRF3 proteins is immobile and organized into stable nanodomains and a much smaller amount is found in a mobile fraction.

**Fig. 4:**
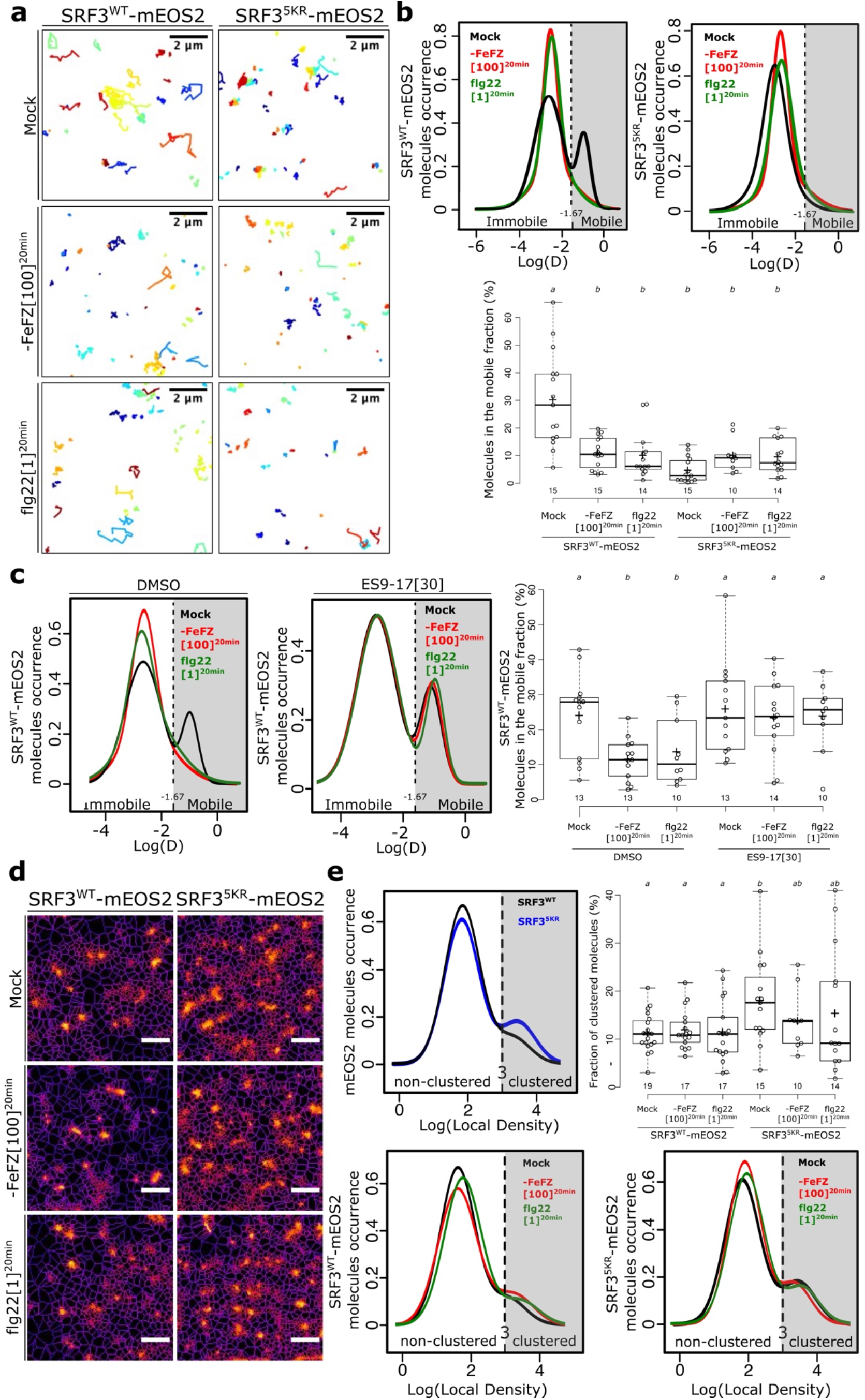
Ubiquitination drives SRF3 nano-organization at the plasma membrane. **a**, Representative *pUBQ10::SRF3^WT^-mEOS2* (SRF3^WT^-mEOS2, left) and *pUBQ10::SRF^5KR^-mEOS2* (SRF3^5KR^-mEOS2, right) trajectories obtained by sptPALM analyses under mock, −Fe supplemented with Ferrozine (−FeFZ) at 100 μM or flg22 at 1μM for 20 minutes. Scale bar, 2μm. **b**. Upper panel: Distribution of SRF3^WT^-mEOS2 and SRF3^5KR^-mEOS2 molecules according to their apparent diffusion coefficient D obtained by analyzing SptPALM trajectories under mock, −Fe supplemented with Ferrozine (−FeFZ) at 100 μM or flg22 at 1μM for 20 minutes. Lower panel: quantification of the percentage of molecules in the mobile fraction. [two-way ANOVA followed by a post hoc Tukey HSD test, letters indicate statistical differences (p<0.05)]. **c**. Left panel: distribution of *pUBQ10::SRF3^WT^-mEOS2* (SRF3^WT^-mEOS2) molecules according to their apparent diffusion coefficient D obtained by analyzing sptPALM trajectories under mock, −Fe supplemented with Ferrozine (−FeFZ) at 100 μM or flg22 at 1μM in DMSO (left) or ES9-17 (right) at 30 μM for 20 minutes. Right panel: quantification of the percentage of molecules in the mobile fraction. [two-way ANOVA followed by a post hoc Fischer LSD test, letters indicate statistical differences (p<0.05)]. **d**, Live PALM analysis of *pUBQ10::SRF3^WT^-mEOS2* (SRF3^WT^-mEOS2, left) and *pUBQ10::SRF^5KR^-mEOS2* (SRF3^5KR^-mEOS2, right) molecules density by tessellation-based automated segmentation of super resolution images under mock, −Fe supplemented with Ferrozine (−FeFZ) at 100 μM or flg22 at 1μM for 20 minutes. Scale bar, 2μm. **e**, Left and lower panel: distribution of *pUBQ10::SRF3^WT^-mEOS2* (SRF3^WT^-mEOS2) and *pUBQ10::SRF^5KR^-mEOS2* (SRF3^5KR^-mEOS2) molecules according to their local density under mock, −Fe supplemented with Ferrozine (−FeFZ) at 100 μM or flg22 at 1μM for 20 minutes. Upper right panel: quantification of the percentage of clustered molecules. [two-way ANOVA followed by a post hoc Fischer LSD test, letters indicate statistical differences (p<0.05)].

### Ubiquitination and degradation steers SRF3 accumulation in the mobile fraction

We then investigated the impact of low iron and flg22-derived signal perceptions on SRF3 partitioning at the plasma membrane. Upon exposure to either flg22 or low iron, the mobile fraction of SRF3^WT^ was drastically decreased from about 30% to 10%, which led us to hypothesize that SRF3 clathrin-dependent endocytosis triggers this decreases (Figure 4a-b – Supplementary Data 5a). To test this latter hypothesis, we performed Spt-PALM applying the clathrin specific inhibitor ES9-17. We observed no decrease of the mobile fraction upon application of ES9-17 during our treatment with low iron or flg22 compared to control conditions (Fig. 4c and Supplementary Data 5b-c). These data confirmed that upon signals perception, the loss of mobile fraction is induced by an increase of SRF3 endocytosis. Overall, our results indicate that SRF3 is accumulated in two fractions in standard conditions and that the SRF3 mobile fraction is decreased due to endocytosis during signal perception. As we had found that SRF3 degradation upon signals is prevented when SRF3 cannot be ubiquitinated in the SRF3^5KR^ line, we analyzed the diffusion of SRF3^5KR^-mEOS2. This version was mainly confined to the immobile fraction representing 95% of the total population in standard condition as well as in low iron and flg22 treated conditions (Fig. 4a-b and Supplementary Data 5d). This result indicated that non-ubiquitinated SRF3 is mainly localized in the immobile fraction and that SRF3 ubiquitination is critical to sustain the mobile fraction. Altogether, we concluded that the mobile fraction is where SRF3 ubiquitination-dependent endocytosis is mediated upon low iron and flg22 treatment. We then decided to investigate the effect of signal perception and ubiquitination on the SRF3 immobile fraction using a cluster analysis approach. While treatments did neither affect cluster density or size in SRF3^WT^and SRF3^5KR^, we observed a general increase and decrease of cluster density and size, respectively, between these two SRF3 versions (Fig. 4d-e and Supplementary Data 6). This shows that while ubiquitination controls the number and the size of clustered SRF3, the presence of signals has no impact on these parameters. Taken together, our results show that SRF3 ubiquitination-dependent endocytosis upon low iron or flg22 signals is mainly regulated by the mobile fraction but not by the immobile nanodomains.

### The distinct low iron and flg22 triggered responses are specified by SRF3 ubiquitination

Since we had previously demonstrated that SRF3 is at the nexus of signaling pathways that are triggered in response to low iron levels and bacterial elicitation by flg 22^27^, we wondered whether SRF3 degradation is required to mediate signaling upon both signals. We first dissected whether SRF3 regulates low iron and flg22 root growth through a common or through independent signaling pathways. To do so, we compared the root growth rate of WT, *srf3-3* and *pSRF3::SRF3^WT^-mCITRINE* in *srf3 (SRF3^WT-comp^)* lines on low iron, flg22 and low iron in combination with flg22. Low iron conditions, supplemented with flg22 elicited a stronger response than observed in each single conditions and mock treatment, suggesting the existence of two distinct root growth signaling pathways (Supplementary Data 7a). In *srf3-3* mutant, we did not detect such a strong decrease and the responsiveness was restored in the complemented *SRF3^WT-comp^* line (Supplementary Data 7a-b). From this experiment, we concluded that *SRF3* controls two independent root growth signaling pathways. We then tested whether SRF3 ubiquitination can specify those signaling pathways. To do so, we exposed WT, *srf3-3, SRF3^WT-comp^* and *pSRF3::SRF3^5KR^-mCITRINE* in *srf3-3* (*SRF3^5KR-comp^*, Supplementary Data 7b) to either low iron levels or flg22 treatment. We monitored early and late root growth responses, which correspond to the analysis of the root growth in these conditions for the first 12 hours or 3 days, respectively. We then measured root growth and then normalized root growth in treatment conditions to the respective standard condition. On low iron medium, roots of *srf3-3* and *SRF3^5KR-comp^* responded less compared to WT and *SRF3^WT-comp^* suggesting that SRF3 ubiquitination-mediated degradation is required to regulate the root growth response to low iron (Fig. 5a-h and Supplementary Data 7c). Surprisingly, the early and late root growth responses to flg22 were rescued in *SRF3^5KR-comp^* plants compared to *srf3-3* plants, showing that SRF3 ubiquitination is not required to mediate root growth arrest in response to flg22 (Fig. 5a-h and Supplementary Data 7c). To corroborate this finding, we calculated the late root growth response to low iron and flg22 in WT and the overexpressing lines *pUBQ10::SRF3^WT^-mCITRINE (SRF3^WT^-OX), pUBQ10::SRF3^5KR^-mCITRINE (SRF3^5KR^-OX)*. In accordance with our previous findings, *SRF3^WT^-OX* was more sensitive to low iron levels^27^, but *SRF3^5KR^-OX* remained completely insensitive compared to WT, arguing that SRF3 ubiquitination is critical to mediate root growth arrest under low iron levels (Supplementary Data 7d). However, we could not detect differences between the late root growth responses of *SRF3^WT^-OX, SRF3^5KR^-OX* and WT under flg22 (Supplementary Data 7e). These results suggest that SRF3 ubiquitination-mediated endocytosis is critical to regulate root growth under low iron levels but not under flg22. Taken together, our results indicated that SRF3 sits at the nexus of two independent root growth signaling pathways, one being specific to low iron which is elicited when SRF3 is ubiquitinated and the other one specific to flg22 signals when SRF3 is non-ubiquitinated.

**Fig. 5:**
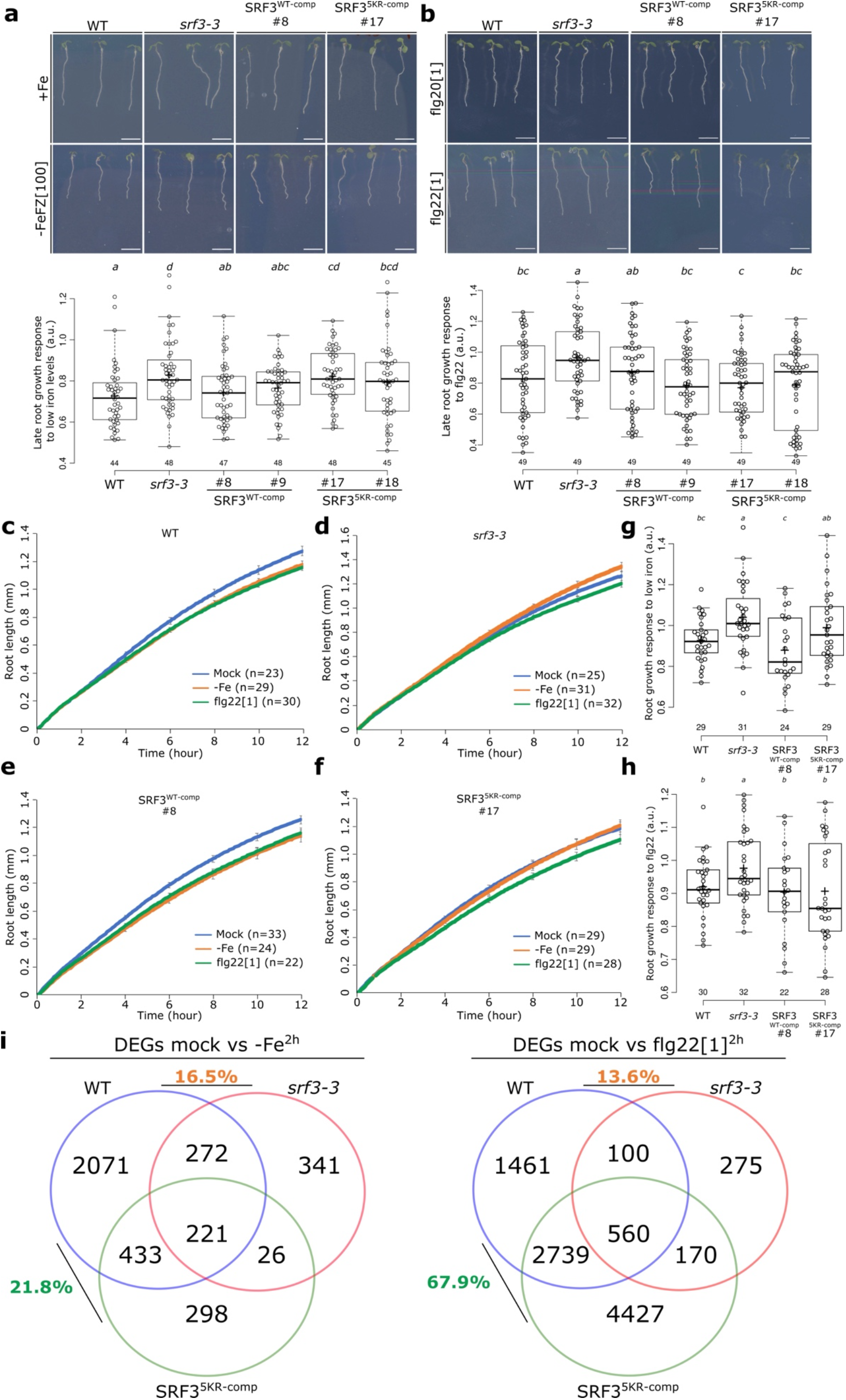
SRF3 ubiquitination determines signaling outputs. **a**, Upper panel: representative images of 8-day old seedlings of WT, *srf3-3, pSRF3::SRF3^WT^-mCITRINE* (*SRF3^WT-comp^*) and *pSRF3::SRF3^5KR^-mCITRINE* (*SRF3^5KR-comp^*) grown under standard conditions for 5 days and then transferred for 3 days to iron sufficient media (+Fe) or to low iron medium supplemented with Ferrozine at 100μM (−FeFZ[100]). Lower panel: quantification of the late root growth response. Scale bar, 1cm [twoway ANOVA followed by a post hoc Fischer LSD test, letters indicate statistical differences (p < 0.05)]. **b**, Upper panel: representative images of 8-day old seedlings of WT, *srf3-3*, *pSRF3::SRF3^WT^-mCITRINE* (*SRF3^WT-comp^*) and *pSRF3::SRF3^5KR^-mCITRINE* (*SRF3^5KR-comp^*) grown under standard conditions for 5 days and then transferred for 3 days to media supplemented with flg20 at 1μM or flg22 at 1μM. Lower panel: quantification of the late root growth response. Numbers indicate independent insertion lines. Scale bar, 1cm. [two-way ANOVA followed by a post hoc Fischer LSD test, letters indicate statistical differences (p<0.05)]. **c-f**, Graph showing time-lapse analysis for 12 h of the root length of WT (**c**), *srf3-3* (**d**), *pSRF3::SRF3^WT^-mCITRINE* (*SRF3^WT-comp^*, **e**) and *pSRF3::SRF3^5KR^-mCITRINE* (*SRF3^5KR-comp^*, **f**) under mock, low (−Fe) iron media and flg22. [error bars indicate standard error of the mean (SEM)] and the related quantification of the early root growth response to low iron (**g**) or to flg22 (**h**). [two-way ANOVA followed by a post hoc Fischer HSD T-test (p<0.07) (g) two-way ANOVA followed by a post hoc Fischer HSD T-test (p<0.08) (h), letters indicate statistical differences]. Numbers indicate independent insertion lines. **i**, Venn diagram of differentially expressed genes (DEGs) under low iron levels (−Fe) for 2 hours (left) or under and flg22 at 1 μM for 2 h (right) in WT, *srf3-3*, and *pSRF3::SRF3^5KR^-mCITRINE* (*SRF3^5KR-comp^*).

To further test this model on a genome-wide scale, we performed an RNAseq analysis in WT, *srf3-3*, and *SRF3^5KR-comp^* lines. Consistent with our previous findings, *SRF3* is critical to regulate transcriptional responses to low iron and flg22 after 2 hours since 16.4% and 13.6% of genes were differentially expressed (DEG) in *srf3-3* compared to WT (Fig. 5i and Supplementary Table 1)^27^. Performing the same experiment with the *SRF3^5KR-comp^* line revealed that while the introgression of *SRF3^5KR^* in *srf3-3* was not able to complement the vast majority of the transcriptome changes of the *srf3* mutant under low iron (in *srf3-3* 16.4% of DEGs overlapped with WT, in *SRF3^5KR^* 21.8 %), it was sufficient to complement a large fraction under flg22 (in *srf3* 13.6% of DEGs overlapped with WT, in *SRF3^5KR^* 67.9 %) (Fig. 5i and Supplementary Table 1). Altogether these results show that depending on the signal perceived, SRF3 ubiquitination status specifies the signaling output since SRF3 ubiquitination is required to mediate low iron responses while non-ubiquitinated SRF3 is sufficient to drive response under bacterial elicitation with flg22.

### Flg22-dependent phosphorylation on Serine 452 leads to SRF3 enrichment in the immobile fraction and prevents low iron signaling

To further test our model that SRF3 is specifically signaling from the immobile fraction upon flg22 elicitation, we analyzed a constitutive active SRF3 version which phosphomimics the flg22-induced SRF3 phosphorylation on serine (S) 452^43^. To do so, we mutated this residue into an aspartic acid (D) and fused it to mEOS2 driven it by the *UBIQUITIN10* promotor (SRF3^S452D^) to then perform Spt-PALM. This experiment revealed that SRF3^S452D^ accumulates preferentially in the immobile fraction compared to the SRF3^WT^ highlighting that SRF3 might specifically signal from the immobile fraction upon flg22-induced phosphorylation, thereby mimicking SRF3^5KR^ (Fig. 6a – Supplementary Data 8a). We then set out to characterize whether SRF3^S452D^ affects SRF3 degradation. Western blots using SRF3^WT^ and SRF3^S452D^ lines in which the SRF3 versions were overexpressed at similar levels revealed that SRF3^S452D^ accumulated more than the WT version in root tissues, similarly to SRF3^5KR^ in basal conditions (Supplementary Data 8b-d). Upon treatment with low iron or flg22, SRF3^S452D^ was not removed from the plasma membrane and accumulated in intracellular compartments in the root transition/elongation zone compared to SRF3^WT^ (Fig. 6b). We then wondered whether this SRF3 phosphomimic version displayed an altered ubiquitination status. Co-IP using ubiquitin antibody revealed no change of the ubiquitination status showing that SRF3 can be ubiquitinated in the immobile fraction while signaling (Supplementary Data 8e). Taken together, these results showed that upon elicitation with flg22 SRF3 phosphorylation on Serine 452 prevents its degradation but not its ubiquitination. This further demonstrates that SRF3 needs to be localized in the mobile fraction and ubiquitinated to be degraded. Consequently, the SRF3 fraction that is degraded upon flg22 signal perception (Fig. 6b) might either have not being phosphorylated or unphosphorylated on this residue or must have other post-translational modifications to be degraded. Finally, we tested whether S452D affects SRF3’s signaling role by assessing the late root growth response to low iron and flg22. Like SRF3^5KR^, the root growth response of *SRF3^S452D^* overexpressing lines of (*UBQ10::SRF3^S452D^-mCITRINE, SRF3^S452D^-OX*) remained insensitive to low iron but restored sensitivity to flg22 compared to the WT (Figure Supplement 8F-G). We then utilized a complementation approach expressing *pSRF3::SRF3^WT^-mCITRINE* in *srf3-3* (*SRF3^WT-comp^*) or *pSRF3::SRF3^S452D^-mCITRINE* in *srf3-3* (*SRF3 ^S452D-comp^*) to test whether like SRF3^5KR^, SRF3^S452D^ does not signal under low iron. As expected, this version did not complement the *srf3-3* root growth phenotype under low iron (Fig. 6c and Supplementary Data 8h-i), showing that SRF3 needs to be localized in the mobile fraction and ubiquitinated to signal in this condition. To sum up, these results indicate that SRF3 flg22-dependent signaling occurs in the immobile fraction. However, the SRF3^S452D^ version that is enriched in the immobile fraction does not signal under low iron levels and according to our previous findings it is not degraded upon signal perception. Yet, this phosphomimic version is still ubiquitinated. This supports a model where SRF3 could be ubiquitinated in the immobile fraction but needs to reach the mobile fraction to mediate its endocytosis and trigger low iron signaling.

**Fig. 6:**
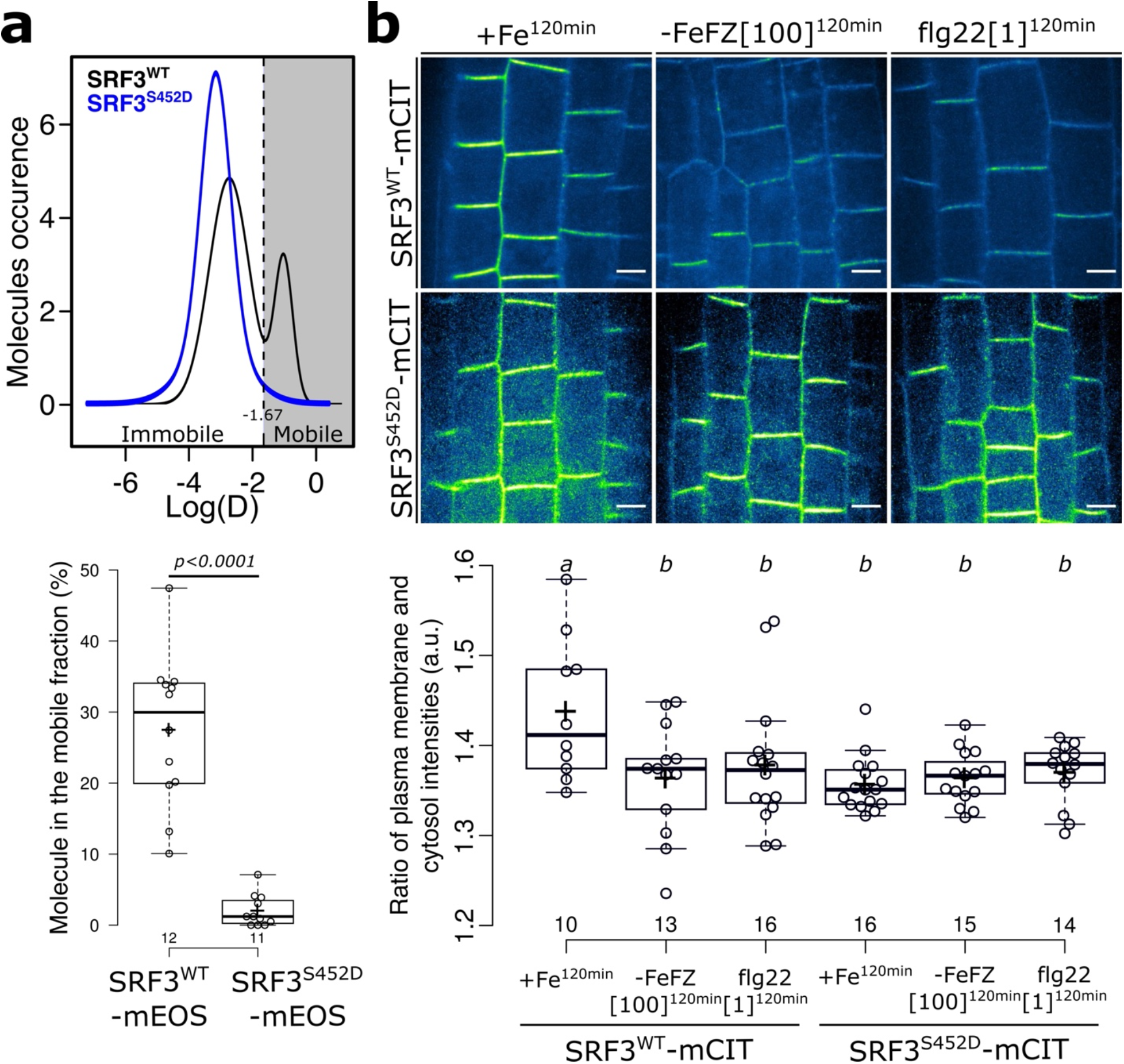
Phosphomimic version of flg22-triggered SRF3 phosphorylation accumulates in the immobile fraction and restricts its degradation upon low iron and flg22. **a**, Upper panel: distribution of *pUBQ10::SRF3^WT^-mEOS2* (SRF3^WT^-mEOS2) and *pUBQ10::SRF^S452D^-mEOS2* (SRF3^S452D^-mEOS2) molecules according to their apparent diffusion coefficient D obtained by analyzing sptPALM trajectories under standard conditions. Lower panel: quantification of the percentage of molecules in the mobile fraction. [Independent two-way Student’s T-test (p<0.05)]. **b**, Upper panel: confocal images of root epidermal cells of 5-day old seedlings expressing *pUBQ10::SRF3^WT^-mCITRINE*, (SRF3^WT^-mCIT) *pUBQ10::SRF3^S452D^-mCITRINE* (SRF3^S452D^-mCIT) under mock, −Fe supplemented with Ferrozine (−FeFZ) at 100 μM or flg22 at 1μM for 120 minutes. Lower panel: quantification of SRF3 plasma membrane mean gray value (a.u.). Scale bar, 5μm. [two-way ANOVA followed by a post hoc Tukey HSD T-test, letters indicate statistical differences (p<0.05)].

## DISCUSSION

Root growth responses to low iron environments and flg22 exposure are linked through the LRR receptor kinase SRF3. Our previous study had shown that SRF3 gets removed from the plasma-membrane in response to these cues^27^. Here, we found that SRF3 is endocytosed by clathrin-dependent endocytosis upon the perception of both signals and that ubiquitination is required for this process. This bears similarities to several other SRF family members that have been predicted to be also ubiquitinated at lysine residues in the juxta membrane domain ^39^. Among them, SRF9/SUB is ubiquitinated and recognized by the clathrin-dependent endocytosis machinery to be addressed to the vacuole for degradation and regulate downstream signaling^38^. Like in other eukaryotic systems, there are alternatives to clathrin-dependent endocytosis^44^. In our study, we did not address this possibility as our experiments using the clathrin specific inhibitor ES9-17 or clathrin heavy chain 1 and 2 (*chc*) mutants indicate that an alternative route might not be relevant for SRF3 levels under low iron and flg22 elicitation (Figure 2c-e).

Super resolution microscopy illustrated that SRF3 segregates into two fractions, the immobile fraction that is organized into nanodomains, and the mobile fraction. Importantly, this nano-organization is required for the specificity of SRF3 dependent signaling and relies on its ubiquitination status. The immobile nanoclustered SRF3 fraction which is non-ubiquitinated is sufficient to mediate flg22 elicited signaling, while ubiquitinated SRF3 that is enriched in the mobile fraction mediates signaling in response to low iron environments. We therefore propose a model in which distinct SRF3 fractions that are defined by their ubiquitination-dependent nanoorganization specifies distinct signaling outputs. This model would explain how a single receptor can mediate different responses based on its post-translational regulation. Additionally, this finding further highlights a new regulatory role of ubiquitination. While being one of the most studied post translational modifications regulating stability, interactions or activity of cellular regulators^45^, we shed light on a new function of ubiquitination in protein organization at the plasma membrane in eukaryotes.

Using super resolution Spt-PALM microscopy, we found evidence that the mobile fraction of SRF3 needs to be ubiquitinated to be sustained, and that upon signals perception this fraction decreases by endocytosis. Phosphorylation of SRF3 might play an important role in this, as the flg22-dependent phosphomimic SRF3 version is enriched in the immobile fraction and while it still can be ubiquitinated, it is not endocytosed upon signal perception. This evidence strongly suggests that SRF3 might be endocytosed in an ubiquitination-dependent manner from the mobile fraction. To support this hypothesis, super resolution imaging on CLC2 revealed that clathrin accumulates in the mobile fraction with up to 30% highlighting that clathrin is present in this fraction^46^. Additionally, the last step of clathrin-dependent endocytosis is relying on the dynamin for clathrin coated tip fission from the plasma membrane. It has been shown that DRP1E has a high diffusion rate further supporting that clathrin-associated cargo proteins might have a high diffusion rate prior being internalized^47^. However, no direct experiments support this model. We therefore cannot exclude a model in which SRF3 is recognized by clathrin adapters when ubiquitinated in the mobile fraction corresponding to the assembly phase, and then the cargo internalization corresponding to the maturation phase occurs in the immobile nanodomains. However, this alternative model does imply a dynamic and interdependent exchange of molecules between the two fractions. While this would be in accordance with our observation about the decrease of nanodomain size and the increase of cluster density when SRF3 ubiquitination is prevented, this interchange of molecules could be independent of the endocytosis process. Overall, therefore both models can explain our observations and future studies are needed to address in detail from which plasma membrane region SRF3 is endocytosed when ubiquitinated.

SRF3 is known to modulate two connected environmental cues, low iron levels and bacterial presence. Accordingly, there is a significant overlap of early responses to low iron and flg22 which is SRF3-dependent^27^. So, what is the relevance of this immediate link between iron and immunity? A decrease in local iron concentrations might due to the growth of other organisms in close vicinity^48^. In particular, bacteria can secrete siderophores to scavenge iron very effectively^48^. A lack of environmental iron therefore could be informative of a potential threat and the activation of defense responses to prime the plants against an eventual bacterial attack to reduce the risk in case of an infection. Thus, in this scenario plants must be able to coordinate both signaling pathways. We provide evidence that SRF3 is central in this coordination and can adjust one or the other pathway depending on the phosphorylation and ubiquitination status that drives its nano-organization at the plasma membrane. Moreover, iron has been shown to be a keystone for host defense and bacterial virulence in plants and mammals^26^. Making iron unavailable to pathogens to restrict their growth is a major part of nutritional immunity and has been proposed to be the first line of the host defense^26,29^. Previously, we have found that SRF3 similarly to the mammalian iron receptor Tfr, regulates immune system and cellular iron availability ^27^. This further illustrates that rapid sensing of bacterial signals and coordination of iron homeostasis is central to mounting an efficient defense response. Since the lack of iron is detrimental to living organisms, the iron homeostasis needs to be tightly balanced upon bacterial presence. Here, we propose that SRF3 through its dual role fine tune those responses to properly adjust iron homeostasis and defense responses. The cellular mechanism that we have uncovered illuminates how iron and defense signaling can converge on a single membrane receptor. This allows fine tuning of signaling output through both, its post-translational modification and dependent partitioning at the plasma membrane to prioritize priming plants against an eventual pathogen attack or low iron responses.

## Acknowledgements

We thank all the Busch lab members for critical discussions. Y. Jaillais for sharing cloning materials. Jiří Friml for providing, *UBQ10::XVE-AUXILIN-LIKE2* and *CLC2::CLC2-GFP*. Y. Jaillais for providing, *pFLOT4::FLOT4-GFP*. We thank, T. Zhang, Rebecca Gilson and the Salk Biophotonics core team for microscopy advance and assistance in quantification. We thank the Salk peptide synthesis core, especially Jill Meisenhelder. We thank, Yvon Jaillais, Alexandre Martinieres and Vincent Bayle for helpful comments on the manuscript. This study was funded by the National Institute of General Medical Sciences of the National Institutes of Health (grant number R01GM127759 to W. Busch), and start-up funds from the Salk Institute for Biological Studies (W. Busch). M.P. Platre was supported by a long-term postdoctoral fellowship (LT000340/2019 L) by the Human Frontier Science Program Organization.

## Author contributions

M.P.P was responsible of all experiments described in the manuscript. M. F.G. assisted and performed Spt-PALM, confocal imaging and quantification of SRF3 plasma membrane levels and root growth experiments. L. B. generated and selected all the transgenic materials and perform western blot and Co-IP. M.C assisted and performed biochemistry experiments and S.B.S. provided material and did a critical analysis of the study. M.P.P. and J.N. performed Co-IP with anti-Ub and western blot. L.Z. analyzed the RNAseq. M.P.P. conceived the study and designed experiments. W.B. provided funding, resources, and supervision. M.P.P. and W.B. wrote the manuscript, and all the authors discussed the results and commented on the manuscript.

## Declaration of interest

The authors declare no conflict of interest.

Data availability. Source data are provided: https://github.com/mplatre/SRF3_Ubiquitination. Raw reads used for the RNAseq are available at NCBI under accession number: PRJNA885223.

Code availability. The code used in this study are provided: https://github.com/mplatre/SRF3_Ubiquitination

## Methods

### Plant materials and growth conditions

For surface sterilization, *Arabidopsis thaliana* seeds that had been produced under uniform growth conditions and were placed for 1 h in opened 1.5-mL Eppendorf tubes in a sealed box containing chlorine gas generated from 130 mL of 10% sodium hypochlorite and 3.5 mL of 37% hydrochloric acid. For stratification, seeds were imbibed in water and stratified in the dark at 4 °C for 2–3 days. Seeds were then put on the surface of Fe-sufficient media described in Gruber et al., 2013, using 12-cm x 12-cm square plates^1^. The Gruber et al., 2013 media contains, 750 μM of MgSO_4_-7H_2_O, 625 μM of KH_2_PO_4_, 1000 μM of NH_4_NO_3_, 9400 μM of KNO_3_, 1500 μM of CaCL_2_-2H_2_O, 0.055 μM of CoCL_2_-6H_2_O, 0.053 μM of CuCl_2_-2H_2_O, 50 μM of H_3_BO_3_, 2.5 μM of KI, 50 μM of MnCl_2_-4H_2_O, 0.52 μM of Na_2_MoO_4_-2H_2_O, 15 μM of ZnCl_2_, 75 μM of Na-Fe-EDTA (sufficient media) or 0 μM of Na-Fe-EDTA (low iron media), 1000 μM of MES adjusted to pH 5.5 with KOH. The *srf3-3, chc1-1, chc2-3, XVE>>AX2*, mutant lines are in Col-0 background and were described and characterized^2–5^. The reporter lines, *pCLC2::CLC2-GFP, pSRF3::SRF3-GFP (Ler), pSRF3::SRF3-mCITRINE, pUBQ10::SRF3-mCITRINE, pUBQ10::SRF3-2xmCHERRY, 2×35s::lti6b-GFP, pUBQ10::mCHERRY-VTI12, pUBQ10::mCHERRY-RabF2a, pVHA-A3::VHA-A3-RFP* are in Col-0 background and were described and characterized^3,6–9^. Plants were grown in long day conditions (16/8h) in walk in growth chambers at 21°C, 50uM light intensity, 60% humidity. During nighttime, temperature was decreased to 15°C.

### Plant transformation and selection

Each construct (see below: “Construction of plant transformation vectors (destination vectors) and plant transformation”), was transformed into the C58 GV3101 *Agrobacterium tumefaciens* strain and selected on YEB media (5g/L beef extract; 1g/L yeast extract; 5g/L peptone; 5g/L sucrose; 15g/L bactoagar; pH 7.2) supplemented with antibiotics (Spectinomycin, Gentamycin). After two days of growth at 28C, bacteria were collected using a single-use cell scraper, re-suspended in about 200 mL of transformation buffer (10mM MgCl2; 5% sucrose; 0.25% silwet) and plants were transformed by the floral dipping method^10^. Plants from the Columbia–0 (Col-0) accession were used for transformation. Primary transformants (T1) were selected *in vitro* on the appropriate antibiotic/herbicide (glufosinate for mCITRINE). Approximately 20 independent T1s were selected for each line. In the T2 generation at least 3 independent transgenic lines were selected using the following criteria when possible: i) good expression level in the root for detection by confocal microscopy, ii) uniform expression pattern, iii) single insertion line (1 sensitive to 3 resistant segregation ratio) and, iv) line with no obvious abnormal developmental phenotypes. Lines were rescreened in T3 using similar criteria as in T2 with the exception that we selected homozygous lines (100% resistant). At this step, we selected one transgenic line for each marker that was used for further analyses and crosses.

### Phenotyping of early root growth responses

Seeds were sowed in +Fe media described in Gruber et al., 2013 and stratified for 2–3 days at 4°C^1^. Five days after planting, about 15 seedlings were transferred to a culture chamber (Lab-Tek, Chambers #1.0 Borosilicate Coverglass System, catalog number: 155361) filled with – Fe or +Fe medium described in Gruber et al., 2013 or +Fe medium containing flg22^1^. Note that the transfer took about 45–60 seconds. Images were acquired every 5 minutes for 12 hours representing 145 images per root in brightfield conditions using a Keyence microscope model BZ-X810 with a BZ NIKON Objective Lens (2X) CFI Plan-Apo Lambda.

### Phenotyping of late root growth responses

Seeds were sowed in +Fe media described in Gruber et al., 2013 and stratified for 2–3 days at 4°C^1^. Five days after planting, 6 plants per genotype were transferred to four 12×12cm plates in a pattern in which the positions of the genotypes were alternating in a block design (top left, top right, bottom left and bottom right). After transfer, the plates were scanned every 24 hours for 3 days using the BRAT software.

### SRF3 Transcript Expression by RT-PCR

Total mRNA was extracted from wild type (WT), SRF3 T-DNA knockout mutant (srf3-3), transgenic lines in Col0 background such as: pUBQ10::SRF3-mCitrine (SRF3^WT^) and pUBQ10::SRF3^5KR^-mCitrine (SRF3^5KR^), pUBQ10::SRF3^S452D^-mCitrine (SRF3^S452D^) pSRF3::SRF3^WT^-mCitrine (SRF3^WT_Comp^), pSRF3::SRF3^5KR^-mCITRINE (SRF3^5KR-Comp^) and pSRF3::SRF3^S452D^-mCITRINE (SRF3^S452D-Comp^) using the Sigma-Aldrich Spectrum Plant Total RNA Kit. cDNA was produced using Applied Biosystems (by Thermo Fisher Scientific) High Capacity cDNA Reverse Transcription Kit. The expression of SRF3 and the ubiquitous TCTP transcripts was tested by PCR using the following primers starting with “RT”: RT_SRF3_F, RT_SRF3_R, RT_TCTP_F and RT-TCTP_R

### Microscopy setup

All imaging experiments except when indicated below, were performed with the following spinning disk confocal microscope set up: inverted Zeiss microscope equipped with a spinning disk module (CSU-X1, Yokogawa, https://www.yokogawa.com) and the prime 95B Scientific CMOS camera (https://www.photometrics.com) using a 40 or 63x Plan-Apochromat objective (numerical aperture 1.4, oil immersion). For colocalization, a 880 Airyscan was used. For SptPALM experiment, the Zeiss Elyra was used. GFP and mCITRINE, were excited with a 488 nm laser (150mW) and fluorescence emission was filtered by a 525/50 nm BrightLine^®^ single-band bandpass filter. mCHERRY was excited with a 561 nm laser (80 mW) and fluorescence emission was filtered by a 609/54 nm BrightLine^®^ single-band bandpass filter (Semrock, http://www.semrock.com/). For quantitative imaging, pictures of root cells were taken with detector settings optimized for low background and no pixel saturation. Care was taken to use similar confocal settings when comparing fluorescence intensity or for quantification.

### TIRF microscopy

Total Internal Reflection Fluorescence (TIRF) Microscopy was conducted using the inverted ONI Nanoimager from Oxford microscope with a 100x Plan-Apochromat objective (numerical aperture 1.50, oil immersion). The optimum critical angle was determined as giving the best signal-to-noise ratio. Images were acquired with about 15% excitation (1W laser power) and taking images every 100ms for 300-time steps.

### SptPALM acquisition

Imaging was performed on a Zeiss Elyra PS1 system with a 100x Apo (numerical aperture 1.46 Oil objective), in TIRF mode. The optimum critical angle was determined as giving the best signal-to-noise ratio. Pixel size was 0.107μm. mEOS was photoconverted using 405nm UV laser power and resulting photoconverted fluorophores were excited using 561nm laser. UV laser power was adjusted to obtain a significant number of tracks without an excessive density to facilitate further analyses (0.01 to 0.08%). 10,000 images time series were recorded at 25 frames per second (40ms exposure time) on a 256 x 256-pixel region of interest. Single molecule detection and tracks reconstruction were made using MTT algorithm and further computational analyses of tracks were made using CBS sptPALM analyzer. For more detail^11^.

### Western Blot

The protein extraction and western blot procedures followed the previously published method with some modifications^12^. About 100 seedlings of 5 days old light grown seedling on mesh of the respective transgenic lines were treated in +Fe, −Fe, or flg22[1] for 4 hours depending on the experiment. The roots were cut and immediately frozen in liquid nitrogen and then were ground with liquid nitrogen and lysed directly with 100 μL1xNuPAGE™ LDS Sample Buffer (Invitrogen™, Cat. NP0008) supplemented with 1x NuPAGE™ Sample Reducing Agent (Invitrogen™, Cat. NP0009) for 15 min on ice. The protein samples were denatured by heating for 10 min at 90 °C and centrifuging at 11,000 × g for 10 min. The supernatant protein samples were separated by NUPAGE 10% Bis-Tris Plus Gel (Invitrogen™, Cat.NW00105BOX) and transferred onto Nitrocellulose membrane using an iBlot 2 Dry Blotting system (Invitrogen™, Cat. IB23001). The GFP signal was detected by western blot with the corresponding antibody (α-GFP-HRP) (Cell Signaling, Cat. No. #4370) 1:2000 diluted in 1% BSA, Merck/Calbiochim, Cat. No.12657.

### Immunoprecipitation

Immunoprecipitation experiments to quantify SRF3 levels required 1–3 g of root material from whole seedlings (12 day old grown on vertically oriented agar plates). Tissues were ground at 4°C and resuspended in 2 mL of ice-cold sucrose buffer (20 mM Tris at pH 8, 0.33 M sucrose, 1 mM EDTA at pH 8, protease inhibitors, 20mM NEM DUB inhibitor). Samples were centrifuged at 5000g for 10 min at 4°C or until the supernatants were clear. The resulting extracts corresponded to the total protein fraction. Total protein was quantified, and equal amounts were loaded on a SDS-PAGE gel. For cellular fractionation, total protein fractions were centrifuged at 20,000g for 45 min at 4°C to pellet microsomes. The resulting supernatant fractions corresponded to the fraction enriched in soluble proteins. The pellet was resuspended in 1 mL of immunoprecipitation buffer (50 mM Tris at pH 8, 150 mM NaCl, 1% Triton X-100) using a 2-mL potter (Wheaton), and was left on a rotating wheel for 30 min at 4°C. Samples were then pelleted at 20,000g for 10 min at 4°C. The supernatant corresponded to the fraction enriched in microsomal-associated proteins. The proteins in each fraction were quantified and immunoprecipitations were performed on 1 mg of proteins using Milteneyi GFP kit. α-P4D1 (Fig. 3c) or α-P4D1-HRP (Fig. S8e) were used to detect general ubiquitination (REF).

### Iron deficiency, flg20 and flg22 treatments

Seeds were sowed in +Fe medium described in Gruber et al., 2013 and stratified for 2–3 days at 4°C^1^. 5-day-old seedlings were treated for 2 or 4 hours with low iron medium or for 2 hours adding 100uM of FerroZine in liquid medium described in Gruber et al., 2013 using 12-well plates^1^. Note that after the addition of FerroZine the pH was adjusted to the same pH=5.7 as the control medium +Fe. However, no change in the pH was detected in agar adjusted with MES as described earlier and in Gruber et al., 2013^59^. For flg22 treatment, Seeds were sowed in +Fe medium described in Gruber et al., 2013 and stratified for 2–3 days at 4°C^1^. 5-day-old seedlings were treated for 2 or 4 hours in iron sufficient media described in Gruber et al., 2013 supplemented with flg22 or flg20 at 1 *μ*M^1^.

### Treatment with BFA, Wm, ES9-17, Dark, and β-Estradiol

Transgenic lines were incubated in 12 well plates containing 25 *μ*M of Brefeldin A (BFA, Sigma, www.sigmaaldrich.com, BFA stock solution at 50 mM in DMSO) in the corresponding liquid Gruber medium (+Fe, −Fe, −FeFZ[100], flg20[1] or flg22[1]) for 2h^1^. Transgenic lines were incubated in 12 well plates containing 30 *μ*M Wortmannin (Sigma, www.sigmaaldrich.com, WM stock solution at 30 mM in DMSO) in the corresponding liquid Gruber media (+Fe, −Fe, −FeFZ[100], flg20[1] or flg22[1]). Transgenic line were placed in the corresponding liquid Gruber media (+Fe, −Fe, −FeFZ[100], flg20[1] or flg22[1]) in 12 well plates and wrapped up in aluminum foil^1^. Transgenic lines were placed in the corresponding agar Gruber media supplemented with β-Estradiol at 2.5 *μ*M for 3 days._In all cases, roots were imaged within a 10-minute time frame window around the indicated time.

### RNAseq

Total RNA was extracted from roots of plants 5 days after germination using RNA protein purification kit (Macherey-Nagel). Next generation sequencing (NGS) libraries were generated using the TruSeq Stranded mRNA library prep kits (Illumina, San Diego, CA, USA). Libraries were sequenced on a HiSeq2500 (Illumina, San Diego, CA, USA) instrument as single read 50bases. NGS analysis was performed using Tophat2 for mapping reads onto the Arabidopsis genome (TAIR10)60,61,HT-seqfor counting reads and EdgeR for quantifying differential expression62.We set a threshold for differentially expressed genes (Fold change (FC) > 2 or FC < −2, FDR < 0.01). Gene ontology analysis was performed using theAgriGOv2onlinetool63. Venn diagrams were generated with the VIB online tool (http://bioinformatics.psb.ugent.be/webtools/Venn/). The plot in Fig. 3a was generated using the online Revigo software (http://revigo.irb.hr/).

## QUANTIFICATION

### Late root growth response

Plates containing seedlings were scanned from days 5 to 9 after transfer to different media in order to acquire images for further quantification of the root growth rate per conditions. Plates were scanned using BRAT software^13^ each day and were stacked together using a macro in Fiji (Macro_Match_Align)^14^. We then calculated the root length for every day per genotype in each condition to evaluate the root growth rate in Fiji using the segmented line. We first calculated the mean of the root growth rate for each day 5 to 6, 6 to 7, 7 to 8. These values were used to calculate the mean of root growth rate for 3 days. Then, we divided the mean of root growth rate for 3 days to a given media for each plant by the mean of root growth rate for 3 days after transfer to the control media for the entire related genotype. This ratio was used as the late root growth response to low iron levels or flg22. Every experiment was repeated twice.

### Early root growth response

Root length for each seedling was recorded for 12 hours taking a picture every 5 minutes and quantified using a Matlab^®^ script (“Root Walker”). From these measurements, we plotted the root length from T0 to T12 after transfer. We then obtained a curve representing the root length after transfer from which we calculated the area under the curve using the following formula “(Root length T1+Root length T2)/2*(Time T2-Time T1)”. Then, we divided the value of the area under the curve after transfer for each plant in a given condition by the area under the curve after transfer to the control media for the entire related genotype. This ratio was used as the early root growth response to low iron levels or flg22. Every experiment has been repeated three times.

### Mean gray value at the plasma membrane

Confocal images were first denoised using an auto local threshold applying the Otsu method with a radius of 25 and a median filter with a radius of 2 in Fiji^14^. To remove every single bright pixel on the generated-binary image the despeckle function was applied. To obtain a plasma membrane skeleton, we detected and removed every intracellular dot using the “Analyze Particles” plugin with the following parameter, size between 0.0001 and 35 000 μm2 and a circularity between 0.18 and 1. Then, we selected and cropped a zone which only showed a proper plasma membrane skeleton. We created a selection from the generated-plasma membrane skeleton and transposed it to the original image to calculate the plasma membrane intensity. This process has been automated in a Macro (Macro_PM/cyt_Intensity). The plasma membrane intensity value was then divided by the total intensity of the image to normalize the plasma membrane intensity. An average of 45 cells were used for quantification per root. Every experiment was repeated three times.

### Normalized intracellular intensity

Confocal images were first denoised using an auto local threshold applying the Otsu method with a radius of 25 and a median filter with a radius of 2 in Fiji^14^. To remove every single bright pixel on the generated-binary image the despeckle function was applied. To obtain plasma membrane skeleton, we detected and removed every intracellular dot using the “Analyze Particles” plugin with the following parameter, size between 0.0001 and 35 000 μm2 and a circularity between 0.18 and 1. Then, we selected and cropped a zone which only showed a proper plasma membrane skeleton. We created a selection from the generated-plasma membrane skeleton and transposed it to the original image and removed the plasma membrane from this image and calculated the mean gray value. This mean gray value was used as the cytosol mean gray value and was then divided by the total intensity of the image to normalize the cytosol intensity and obtain the normalized intracellular intensity. This process has been automated in a Macro (Macro_PM/Cyt_Intensity). The plasma membrane intensity value was then divided by the total intensity of the image to normalize the plasma membrane intensity. An average of 45 cells were used for quantification per root. Every experiment was repeated three times.

### Ratio of plasma membrane and cytosol intensity

Confocal images were first denoised using an auto local threshold applying the Otsu method with a radius of 25 and a median filter with a radius of 2 in Fiji^14^. To remove every single bright pixel on the generated-binary image the despeckle function was applied. To obtain plasma membrane skeleton, we detected and removed every intracellular dot using the “Analyze Particles” plugin with the following parameter, size between 0.0001 and 35 000 μm2 and a circularity between 0.18 and 1. Then, we selected and cropped a zone which only showed a proper plasma membrane skeleton. We created a selection from the generated-plasma membrane skeleton and transposed it to the original image. First, we calculated the mean gray value of this zone, corresponding to the plasma membrane mean gray value. Then on the original image the same zone has been removed from and the mean gray value was calculated to obtain the cytosol mean gray value. The plasma membrane mean gray value was then divided by cytosol mean gray value to normalize to obtain the ration of the plasma membrane and cytosol intensities. This process has been automated in a Macro (Macro_PM/Cyt_Intensity). An average of 45 cells were used for quantification per root. Every experiment was repeated three times.

### Ratio of the root tip and differentiated zones intensity

The root tip containing the meristematic and elongation zones excluding the root cap was selected and the mean gray value was recorded using Fiji^14^. Then a zone corresponding to the same size area was selected in the differentiation zone and the mean gray value was recorded. The root mean gray value was divided by the differentiation zone mean gray value to obtain the ratio of the root tip and differentiated zones intensity.

### Size of BFA bodies

BFA body size was quantified on at least 28 roots representing an average of 1204 cells in three independent experiments. Threshold was applied, images harboring less than ten BFA bodies were removed from the analysis as well as images issue from misshapen root cells. Cells containing BFA bodies whose length are larger than width were selected for the analysis. Images were submitted to a NidBlack auto local threshold with a radius of 15. BFA bodies bigger than 5.2 um2 with a circularity between 0.25 and 1 were detected using the “Analyze Particles” plugin of Fiji^14^. The average size of BFA bodies was obtained for each root. Per root an average of 38 BFA bodies was detected representing at least 532 BFA bodies quantified per conditions. All this process has been automated through a macro (BFA_Counter). In each graph relative to the size of BFA body, “n” represents the number of roots used for quantification.

### Number of Wm-positive late endosomes per cell

The number of Wm-positive endosomes showing the typical donut-shape like structure were counted by eyes and divided by the number cells were counted in the image to obtain the number of Wm-positive late endosomes per cell.

### SptPALM data quantifications and representations

For each movie corresponding to one condition/genotype, the natural logarithm of the apparent diffusion coefficient (log(D)) per molecule was extracted from the MTT data using the CBS sptPALM analyzer^11^. For sptPALM analyses, only tracks with at least five successive points were selected and the rest of the molecules was discarded from the analysis. We pooled values from each condition in order to get the log(D) distribution using the R software. The set of molecules which present a log(D) value below −1.67 refers to the immobile group, whereas values above this threshold denote the mobile group. To obtain the percentage of molecules in the immobile group, we divided the number of tracks below −1.67 by the total track number multiplied by one hundred. Then, each condition was analyzed using an R script to automatize the analysis. To statistically test the existence of these two behaviors (mobile and immobile), we analyzed the log(D) distribution per condition using the mclust R package. For each condition, the percentages of data in each of the 2 groups defined by the partition P with the absolute log(D) value (−1.67) are first reported. This value was determined following a mixture model of scale mixtures of skew-normal denoted as FMSMSN. We then fitted two distinct two-component Gaussian mixture models, M0 and M1, using the mclust::Mclust R function. M0 is a model where σ1 ≠σ2 and M1 is a model where σ1 = σ2. For each model, every molecule is assigned to one of the 2 Gaussian components Gi (i □_[1, 2]) according to a MAP (Maximum A Posteriori)-based classification. M1 model was used since it provided more representative results despite of a lower Bayesian information criterion values. The percentage of values assigned to each of the two Gaussian components is reported as well as its 95% confidence interval obtained with a non-parametric bootstrap method, using the mclust::MclustBootstrap R function with default arguments. The classification rate was calculated as the percentage of MAP-classified values concordant with the partition P. Based on the estimation of the model M1, we plotted the density of the Gaussian mixture according to the kernel density estimation (called “occurrence” for more clarity) and was used for representation. The R script can be found under “Script_mCLUST_R_SALK_DIFFUSION”. At least 14 acquisitions from three independent experiments were used. In each graph relative to the percentage of immobile, “n” represents the number of acquisitions used for quantification. Note that one acquisition may represent more than one cell but that different acquisitions are always from different cells. For more details refer to^11^.

### LivePALM, Voronoi tessellation analyses and representation

For each movie corresponding to one condition/genotype, the object localization was extracted from MTT data (33) by CBS sptPALM analyzer^11^. SR-tesseler software was used to reconstruct live PALM images and determine protein local density^15^. Multiple detections of single object were corrected as previously described by Levet et al. 2015^15^. Local density was extracted from Voronoï segmentation of the image as shown on Fig. 4d. We pooled values from each condition in order to get the log(local density) distribution using R software. A first partition P was done; the set of molecules which present a log(local density) value below 3 refers to the immobile group, whereas values above this threshold denote the mobile group. To obtain the percentage of molecules in the immobile group, we divided the number of molecules below 3 by the total molecules number multiplied by one hundred. Then, each condition was analyzed using an R script to automatize the analysis. To statistically test the existence of these two behaviors (mobile and immobile), we analyzed the local density distribution per condition using the mclust R package. For each condition, the percentages of data in each of the 2 groups defined by the partition P with the absolute log(local density) value (3) are first reported. We then fitted two distinct two-component Gaussian mixture models, M0 and M1, using the mclust::Mclust R function. M0 is a model where σ1 ≠ σ2 and M1 is a model where σ1 = σ2. For each model, every molecule is assigned to one of the 2 Gaussian components Gi (i □ [1,2]) according to a MAP (Maximum A Posteriori)-based classification. M1 model was used since it provided more representative results despite of a lower Bayesian information criterion values. The percentage of values assigned to each of the two Gaussian components is reported as well as its 95% confidence interval obtained with a non-parametric bootstrap method, using the mclust::MclustBootstrap R function with default arguments. The classification rate was calculated as the percentage of MAP-classified values concordant with the partition P. Based on the estimation of the model M1, we plotted the density of the Gaussian mixture according to the kernel density estimation (called “occurrence” for more clarity) and used for representation. The R script can be found under “Script_mCLUST_R_SALK_LocalDensity”. At least 24 cells from three independent experiments were used. In each graph relative to the fraction of clustered SRF3 in percentage, “n” represents the number of roots used for quantification.

### Clusters density and size analysis

For each movie corresponding to one condition/genotype, the local density of each molecule was calculated with SR-Tesseler^15^. For cluster detection, a threshold of 20 times the average localization density was applied to identify regions with high local density. “Objects” with at least 5 molecules were identified as “clusters” of molecules and from which the size of each cluster was extracted. n indicates the number of detected clusters. The R script for analysis and representation can be found under “ Script_mCLUST_Cluster_diameter”

## STATISTICS

Each experiment has been repeated independently at least twice, as in every case the same trend has been recorded for independent experiments, the data have been pooled for further statistical analysis. Each sample were subjected to four different normality tests (Jarque-Bera, Lilliefors, Anderson-Darling and Shapiro-Wilk), sample were considered as a Gaussian distribution when at least one test was significant (p<0.05) using Xlstat.

- As a normal distribution was observed a two-ways ANOVA coupled with post hoc Tukey test was performed (p<0.05) using Xlstat.
- As a normal distribution was observed at two-ways ANOVA coupled with post hoc Fisher test was performed (p<0.05, p<0.07, p<0.08 and p<0.1) using Xlstat:
- As a normal distribution was observed an independent two-ways student test was performed (p=0.05) using Xlstat.
- As a normal distribution was not observed at two-ways Kruskal-Wallis coupled with post hoc Steel-Dwass-Critchlow-Fligner procedure was performed (p<0.05) using Xlstat.

The type of test performed in indicated in the figure legend.

## CLONING

### SRF3 mutagenesis

SRF3 mutant impaired for ubiquitination was obtained by phosphorylated primers or site directed mutagenesis using SRF3-cds_p221 as a template. 5’ phosphorylated primers such as SRF3_cDNA_K391392R_p_F and SRF3_cDNA_K391392R_p_R were used to mutate lysines (K) 391 and 392 into arginines (R). Then, lysines (K), 444, 458, 471 were mutated into arginine (R) by site-directed mutagenesis using the following primers. SRF3_K444R_F, SRF3_K444R_R, SRF3_K458R_F, SRF3_K458R_R, SRF3_K471R_F and SRF3_K471R_R. SRF3 phosphomimic mutant impaired on Serine (S) 452 was obtained by site directed mutagenesis using SRF3-cds_p221 as a template using the primer SRF3_S452D_F and SRF3_S452D_R.

### List of pENTR clones

List of pENTR clones:
| Name of entry vector | Source | Reference |
| --- | --- | --- |
| SRF3prom_pDONR4P1R | Platre et al., 2022<sup>3</sup> | N/A |
| UBQ10prom_pDONR4P1R | Jaillais et al., 2011<sup>16</sup> | NASC# N2106315 |
| SRF3<sup>5KR</sup>_P1P2 | This study | N/A |
| SRF3<sup>S452D</sup>_P1P2 | This study | N/A |
| mCITRINE_pDONR2RP3 | Simon et al., 2014<sup>17</sup> | NASC# N2106287 |
| mCITRINE-Ub_pDONR2RP3 | Martins et al., 2015<sup>18</sup> | N/A |
| mCITRINE-Ub-I44A_pDONR2RP3 | Martins et al., 2015<sup>18</sup> | N/A |
| mCITRINE-Ub-7KR_pDONR2RP3 | Martins et al., 2015<sup>18</sup> | N/A |
| mEOS2_pDONR2RP3 | Martiniere et al., 2019<sup>19</sup> | N/A |

### Construction of plant transformation vectors (destination vectors) and plant transformation

Binary destination vectors for plant transformation were obtained using the multisite LR recombination system (life technologies, http://www.thermofisher.com/) using the pB7m34GW (basta resistant)

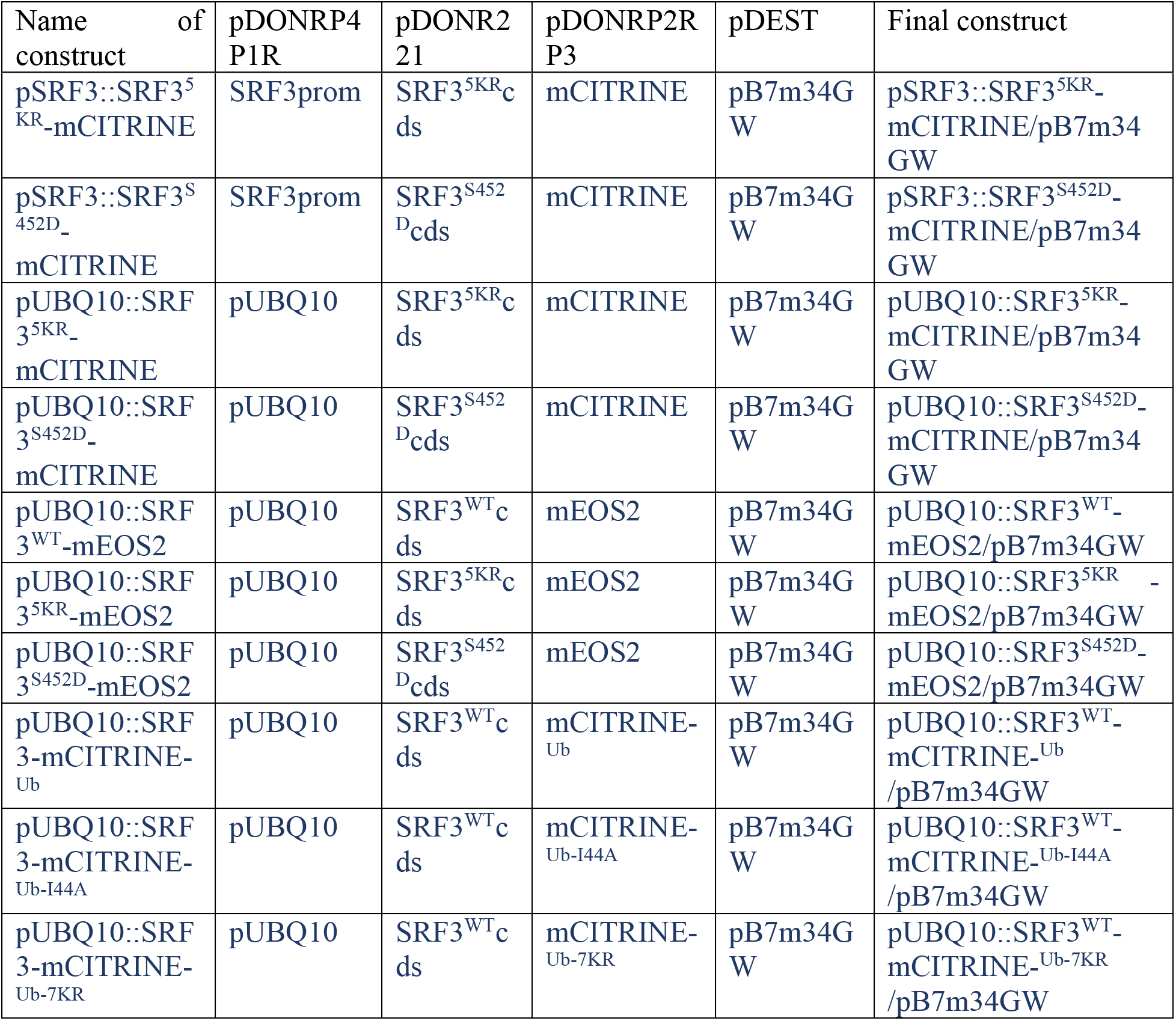

**Primer table:**

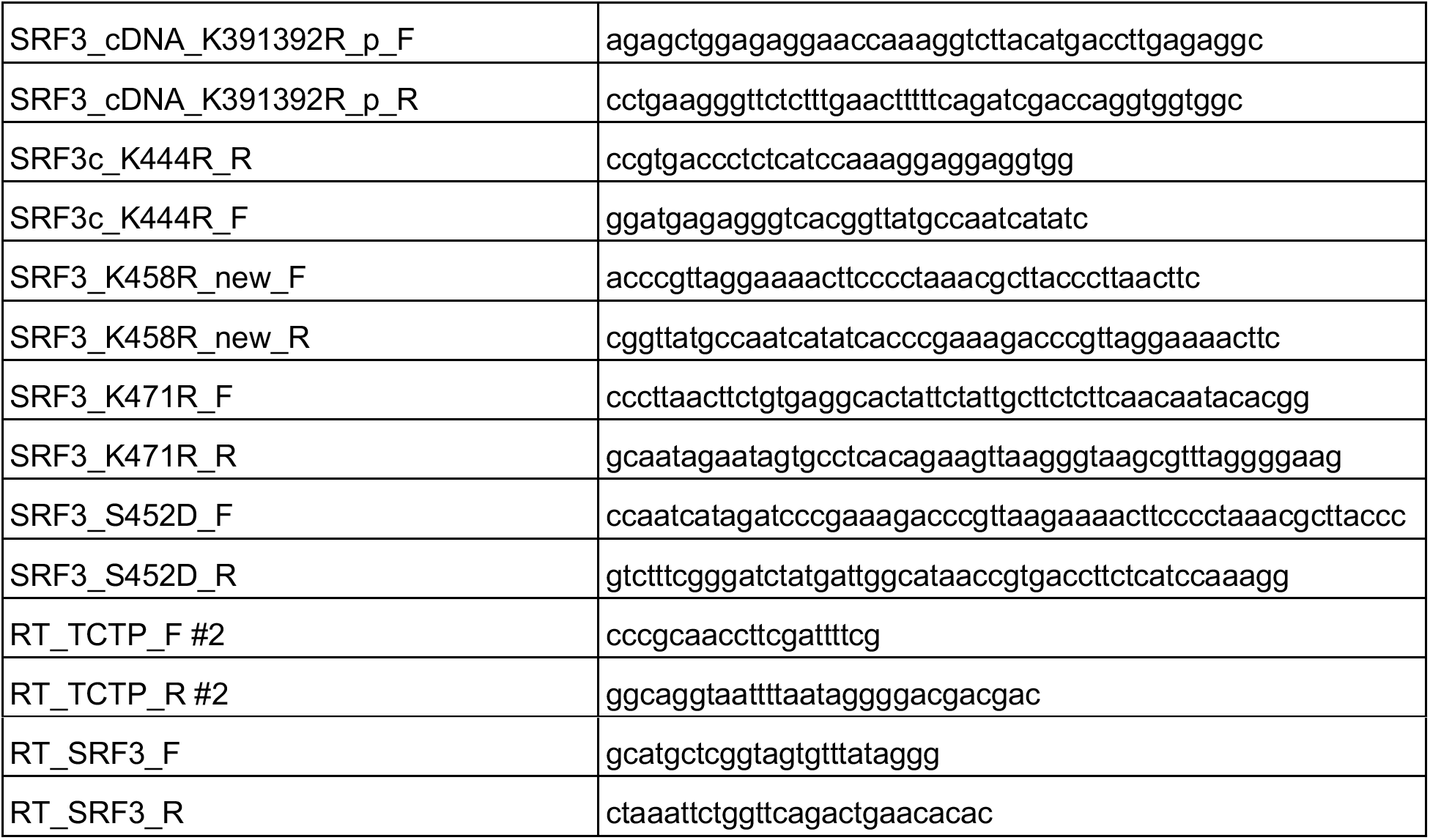

