## Supplementary Figures for "Ubiquitination steers SRF3 plasma membrane nano-organization to specify signaling outputs"

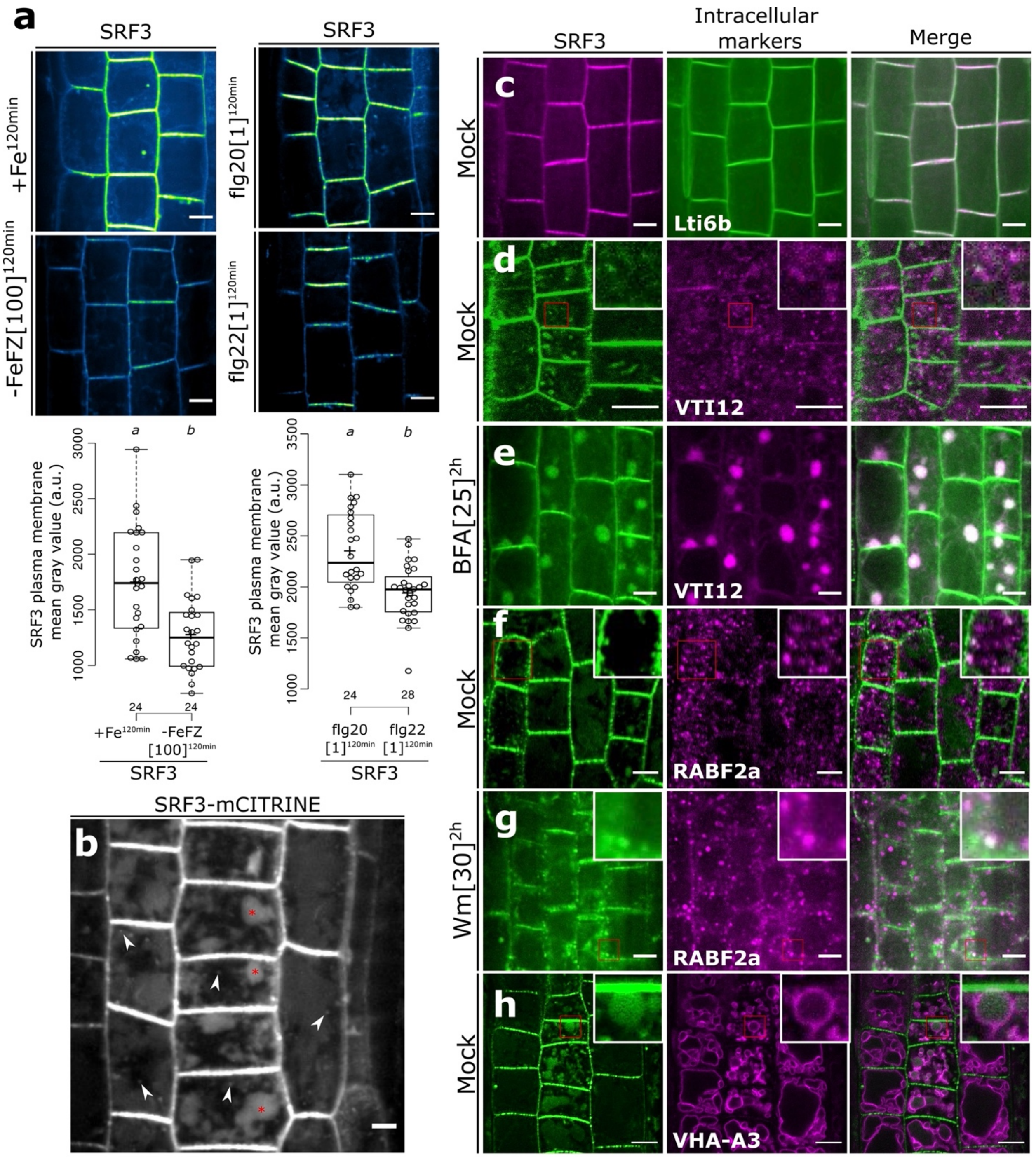

a
SRF3
+Fe120min
-FeFZ[100]120min
flg20[1]120min
flg22[1]120min
SRF3 plasma membrane mean gray value (a.u.)
a
b
24
24
28
+Fe120min
-FeFZ [100]120min
flg20 [1]120min
flg22 [1]120min
b
SRF3-mCITRINE
Intracellular markers
Merge
c
d
e
f
g
h
Mock
BFA[25]2h
Wm[30]2h
Lti6b
VTI12
RABF2a
VHA-A3

**Figure S1: SRF3 is removed from the plasma membrane under low iron and flg22 and accumulates along the endocytosis pathway. a**, Upper panel: confocal images of root epidermal cells of 5-day old seedlings expressing *pUBQ10::SRF3$^{WT}$-mCITRINE* (SRF3) under +Fe, -Fe supplemented with Ferrozine (-FeFZ) at 100 µM for 120 minutes (left) and under flg20 or flg22 at 1µM for 120 minutes (right). Lower panel: quantification of SRF3 plasma membrane mean gray (a.u.). Scale bar, 5µm. [Independent two-way Student's T-test ($p < 0.05$)]. Note that these images and quantification are the same as presented in Figure 2c-d. 3**b**, Confocal images of root epidermal cells of 5-day old seedlings expressing *pUBQ10::SRF3$^{WT}$-mCITRINE* (SRF3-mCITRINE). **c-h**, , Confocal images of root epidermal cells of 5-day old seedlings co-expressing *pUBQ10::SRF3-2xmCHERRY (*SRF3*)* with *p35s::GFP-Lti6b* (Lti6b, **c**), and *pUBQ10::SRF3-mCITRINE (*SRF3*)* with *pUBQ10::VTI12-2xmCHERRY* (VTI12) under mock (**d**) or under BFA at 25µM for 2 hours (**e**), and *pUBQ10::SRF3-mCITRINE (*SRF3*)* with *pUBQ10::RABF2a-2xmCHERRY* (RABF2a) under mock (**f**) or under Wm at 30µM for 2 hours (**g**), and *pUBQ10::SRF3-mCITRINE* (SRF3) with *pVHA-A3::VHA-A3-RFP* (VHA-A3, **h**). Scale bar, 5µm.

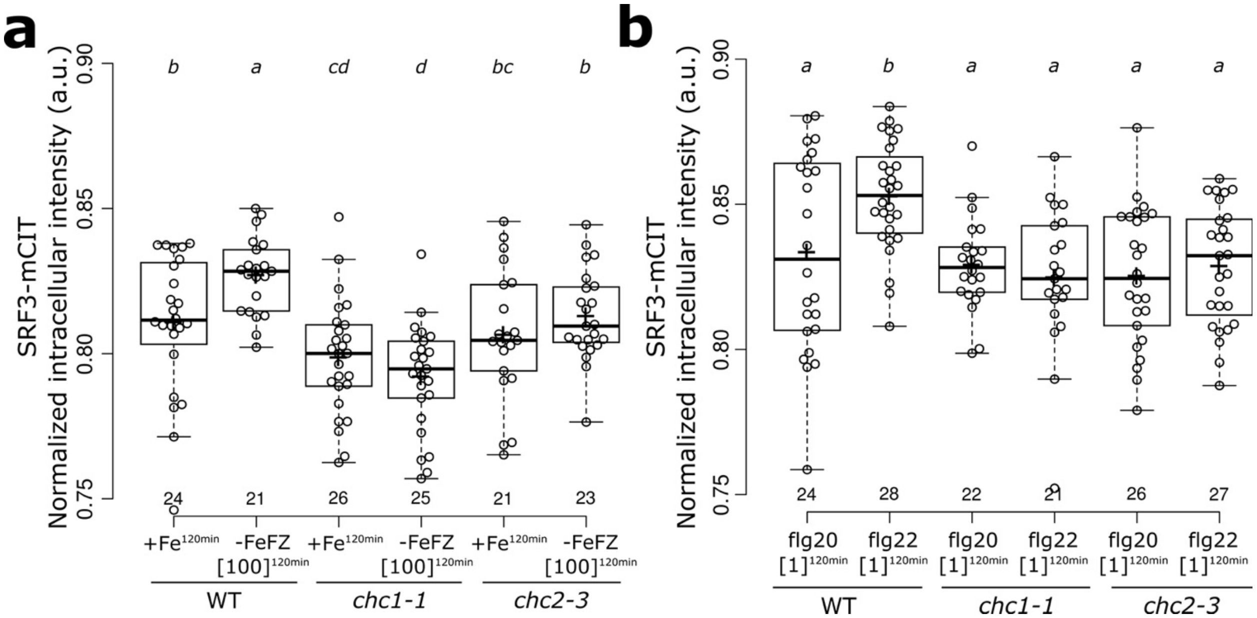


**Figure S2: Clathrin-dependent endocytosis regulates SRF3 enrichment in the intracellular compartments**. **a-b**, Quantification of the normalized intracellular intensity (a.u.) based *on* confocal images of root epidermal cells of 5-day old seedlings expressing *pUBQ10::SRF3-mCITRINE* (SRF3-mCIT) in WT, *chc1-1* or *chc2-3* mutants under +Fe, -Fe supplemented with Ferrozine (-FeFZ) at 100 µM for 120 minutes (**a**) . [two-way ANOVA followed by a post hoc Fischer LSD test, letters indicate statistical differences ($p < 0.05$)] and treated with flg20 or flg22 at 1 µM for 120 minutes (**b**). [two-way ANOVA followed by a post hoc Fischer LSD test, letters indicate statistical differences ($p < 0.05$)].

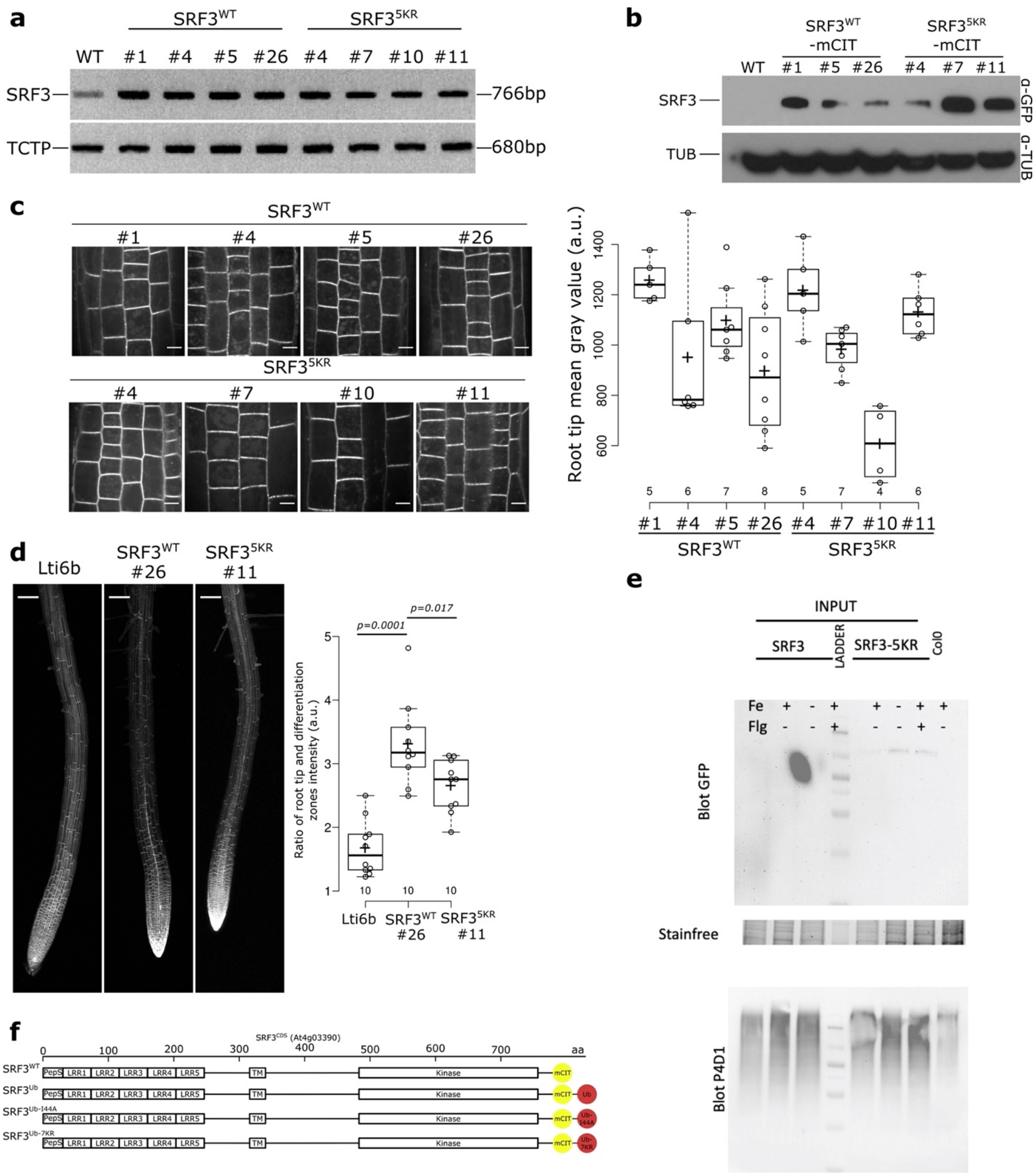

a
SRF3WT
SRF35KR
WT #1 #4 #5 #26 #4 #7 #10 #11
SRF3
766bp
TCTP
680bp
b
SRF3WT -mCIT
SRF35KR -mCIT
WT #1 #5 #26 #4 #7 #11
SRF3
α-GFP
TUB
α-TUB
c
SRF3WT
#1 #4 #5 #26
SRF35KR
#4 #7 #10 #11
Root tip mean gray value (a.u.)
1400 1200 1000 800 600
5 6 7 8 5 7 4 6
#1 #4 #5 #26 #4 #7 #10 #11
SRF3WT
SRF35KR
d
Lti6b
SRF3WT #26
SRF35KR #11
p=0.0001
p=0.017
Ratio of root tip and differentiation zones intensity (a.u.)
5 4 3 2 1
10 10 10
Lti6b
SRF3WT #26
SRF35KR #11
e
INPUT
SRF3
LADDER
SRF3-5KR
Col0
Fe
Flg
Blot GFP
Stainfree
Blot P4D1
f
SRF3CDS (At4g03390)
0 100 200 300 400 500 600 700 aa
SRF3WT
SRF3Ub
SRF3Ub-I44A
SRF3Ub-7KR
PepS LRR1 LRR2 LRR3 LRR4 LRR5 TM Kinase mCIT Ub
Ub-I44A
Ub-7KR

**Figure S3: Lysines 391, 392, 444, 458 and 471 are ubiquitinated on SRF3 and regulate protein levels in root tissues upon low iron and flg22 treatment.** **a**, RT-PCR using SRF3 primers in four different insertion lines of *pUBQ10::SRF3$^{WT}$-mCITRINE* (SRF3$^{WT}$), *pUBQ10::SRF3$^{5KR}$-mCITRINE* (SRF3$^{5KR}$) of 5-day old seedlings. **b**, Western blot on root material of tgree different insertion lines of *pUBQ10::SRF3$^{WT}$-mCITRINE* (SRF3$^{WT}$), *pUBQ10::SRF3$^{5KR}$-mCITRINE* (SRF3$^{5KR}$) of 7-day old seedlings using α-GFP and α-TUB. **c**, Left panel: confocal images of root epidermal cells of 7-day old seedlings expressing *pUBQ10::SRF3$^{WT}$-mCITRINE* (SRF3$^{WT}$). *pUBQ10::SRF3$^{5KR}$-mCITRINE* (SRF3$^{5KR}$ in four different insertions lines. Right panel: quantification of the fluorescence in the root tip. Scale bar, 5µm. Numbers indicate independent insertion lines. Independent two-way Student's T-test ($p < 0.05$)]. **d**, Left panel: confocal images of roots of 7-day old seedlings expressing, *p35s::GFP-Lti6b* (*Lti6b*) or *pUBQ10::SRF3$^{WT}$-mCITRINE* (SRF3$^{WT}$) or *pUBQ10::SRF3$^{5KR}$-mCITRINE* (SRF3$^{5KR}$). Right panel: quantification of the ratio of the root tip and differentiation zone intensity (a.u., right). Scale bar, 50µm. **e**, Western blot of the data presented in Fig. 3b using α-GFP on roots of *pUBQ10::SRF3$^{WT}$-mCITRINE* (*SRF3$^{WT}$*) or *pUBQ10::SRF3$^{5KR}$-mCITRINE* (*SRF3$^{5KR}$*) of 12-day old seedlings under +Fe or flg22 at 1 µM for 120 minutes (left) and below the related co-IP using α-P4D1. **f**, Schematic representation of *pUBQ10::SRF3$^{WT}$-mCITRINE* (SRF3$^{WT}$), *pUBQ10::SRF3-mCITRINE$^{Ub}$* (*SRF3$^{Ub}$*), *pUBQ10::SRF3-mCITRINE$^{Ub\text{-}I44A}$* (*SRF3$^{Ub\text{-}I44A}$*), *pUBQ10::SRF3-mCITRINE$^{Ub\text{-}7KR}$* (*SRF3$^{Ub\text{-}7KR}$*).

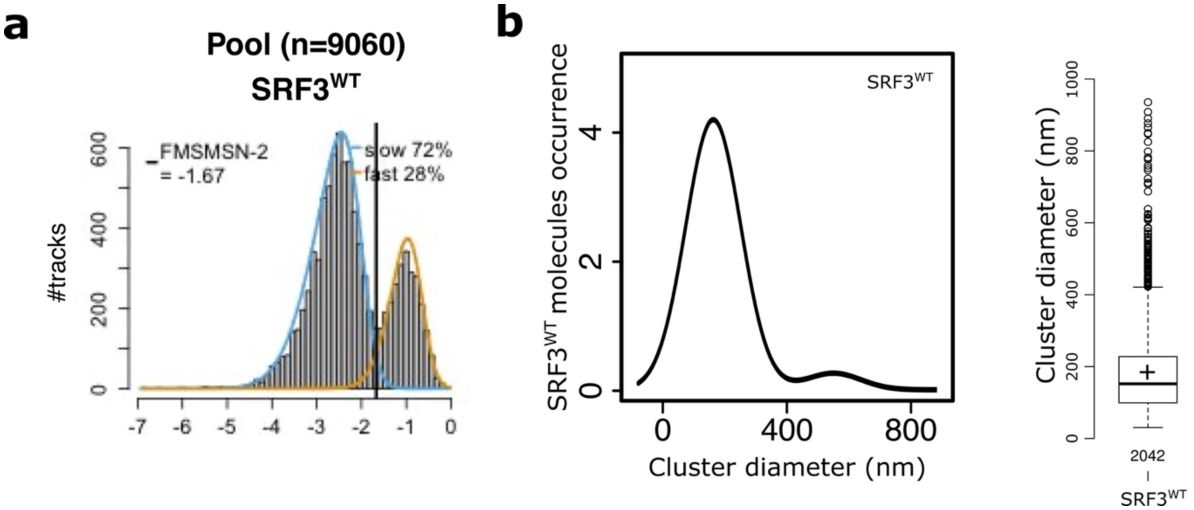


**Figure S4: SRF3 segregates into a mobile fraction and immobile nanoclusters at the plasma membrane. a**, Distribution of *pUBQ10::SRF3$^{WT}$-mEOS2 (*SRF3$^{WT}$-mEOS2) molecules according to their apparent diffusion coefficient D obtained by analyzing sptPALM trajectories under mock condition and separated in two fractions according to a mixture model of scale mixtures of skew-normal. **b**, Left panel: distribution of *pUBQ10::SRF3$^{WT}$-mEOS2 (*SRF3$^{WT}$-mEOS2) molecules according to cluster diameter obtained by analyzing PALM trajectories and clustering analysis according to SR-Tesseler under mock condition. Right panel quantification of the cluster diameter (nm).

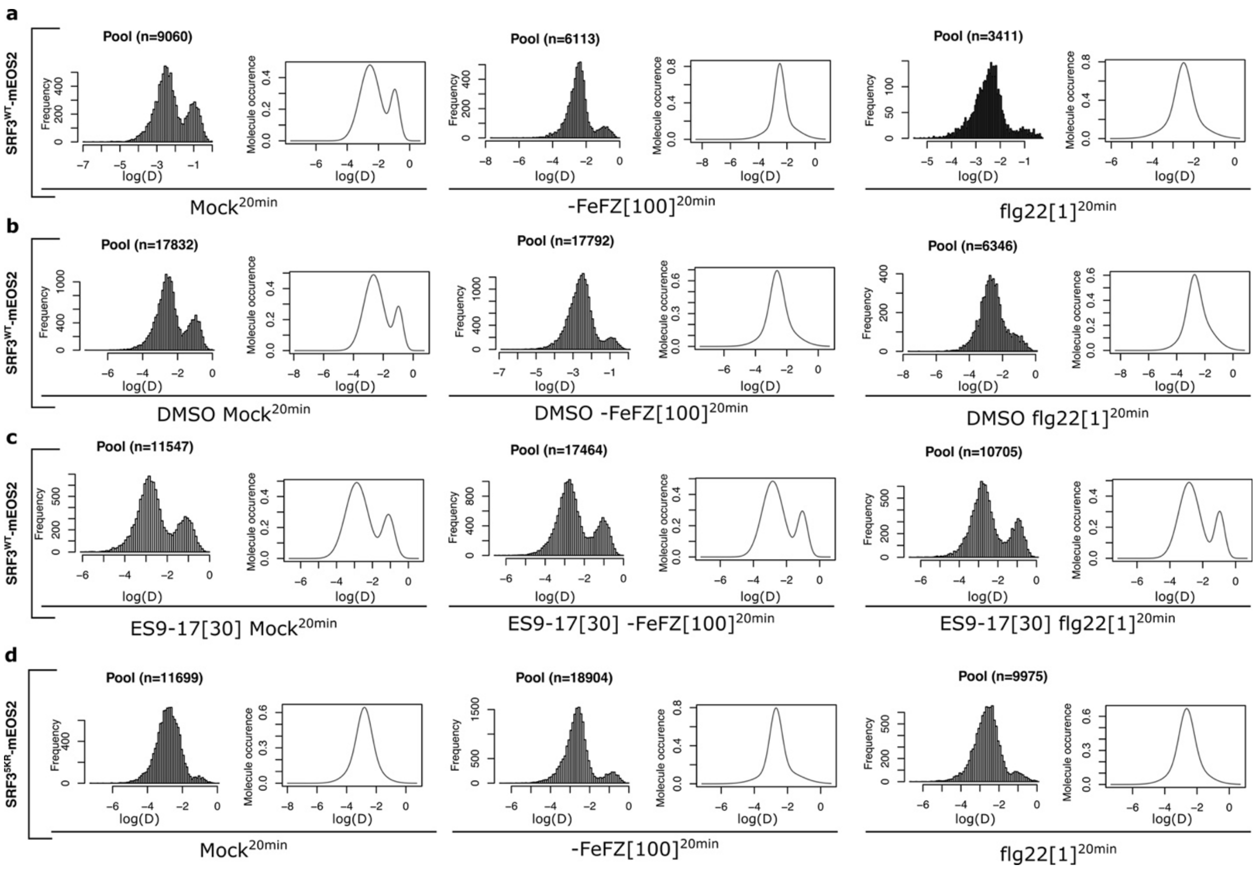
a
SRF3WT-mEOS2
Pool (n=9060)
Pool (n=6113)
Pool (n=3411)
Mock20min
-FeFZ[100]20min
flg22[1]20min
b
SRF3WT-mEOS2
Pool (n=17832)
Pool (n=17792)
Pool (n=6346)
DMSO Mock20min
DMSO -FeFZ[100]20min
DMSO flg22[1]20min
c
SRF3WT-mEOS2
Pool (n=11547)
Pool (n=17464)
Pool (n=10705)
ES9-17[30] Mock20min
ES9-17[30] -FeFZ[100]20min
ES9-17[30] flg22[1]20min
d
SRF35KR-mEOS2
Pool (n=11699)
Pool (n=18904)
Pool (n=9975)
Mock20min
-FeFZ[100]20min
flg22[1]20min
Frequency
Molecule occurence
log(D)

**Figure S5: Ubiquitination-dependent endocytosis regulates SRF3 enrichment in the mobile fraction upon low iron and flg22. a**, Distribution of *pUBQ10::SRF3$^{WT}$-mEOS2 (*SRF3$^{WT}$-mEOS2) molecules according to their apparent diffusion coefficient D obtained by analyzing sptPALM trajectories under mock (left), -Fe supplemented with Ferrozine (-FeFZ) at 100 µM (middle) or flg22 at 1µM (right) for 20 minutes. Smoothed curves represent the distribution of SRF3$^{WT}$-mEOS2 molecules according to their apparent diffusion coefficient (log(D)) obtained by analyzing the frequency plot using the R mClust package. **b**, Distribution of *pUBQ10::SRF3$^{WT}$-mEOS2 (*SRF3$^{WT}$-mEOS2) molecules according to their apparent diffusion coefficient D obtained by analyzing sptPALM trajectories under DMSO with mock (left), -Fe supplemented with Ferrozine (-FeFZ) at 100 µM (middle) or flg22 at 1µM (right) for 20 minutes. Smoothed curves represent the distribution of *pUBQ10::SRF3$^{WT}$-mEOS2 (*SRF3$^{WT}$-mEOS2) molecules according to their apparent diffusion coefficient (log(D)) obtained by analyzing the frequency plot using the R mClust package. **c**, Distribution of *pUBQ10::SRF3$^{WT}$-mEOS2 (*SRF3$^{WT}$-mEOS2) molecules according to their apparent diffusion coefficient D obtained by analyzing sptPALM trajectories under ES9-17 at 30 µM with mock (left), -Fe supplemented with Ferrozine (-FeFZ) at 100 µM (middle) or flg22 at 1µM (right) for 20 minutes. Smoothed curves represent the distribution of SRF3$^{WT}$-mEOS2 molecules according to their apparent diffusion coefficient (log(D)) obtained by analyzing the frequency plot using the R mClust package. **d**, Distribution of *pUBQ10::SRF3$^{5KR}$-mEOS2* (SRF3$^{WT}$-mEOS2) molecules according to their apparent diffusion coefficient D obtained by analyzing sptPALM trajectories under mock (left), -Fe supplemented with Ferrozine (-FeFZ) at 100 µM (middle) or flg22 at 1µM (right) for 20 minutes. Smoothed curves represent the distribution of *pUBQ10::SRF3$^{WT}$-mEOS2 (*SRF3$^{WT}$-mEOS2) molecules according to their apparent diffusion coefficient (log(D)) obtained by analyzing the frequency plot using the R mClust package.

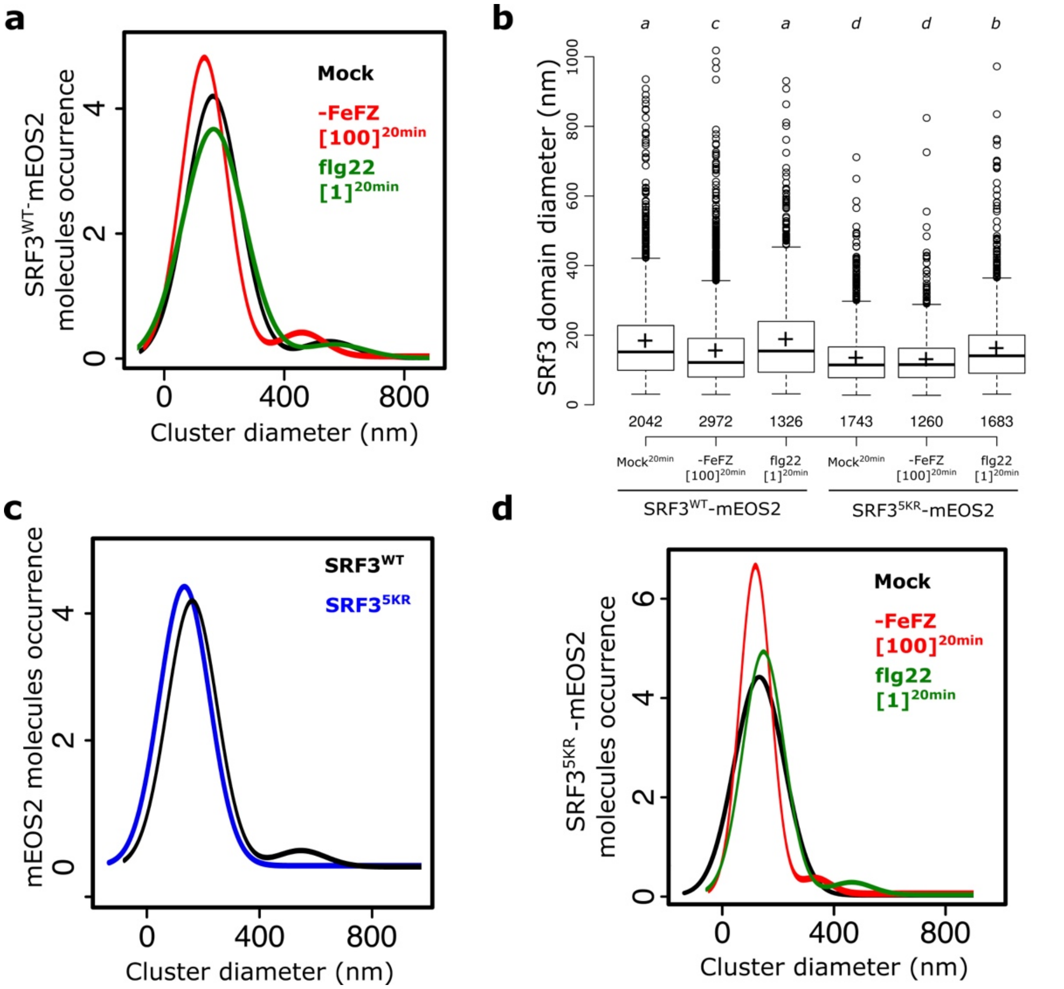


**Figure S6: SRF3 ubiquitination modulates nanoclusters size.** **a**, Live PALM analysis of *pUBQ10::SRF3$^{WT}$-mEOS2* (SRF3$^{WT}$-mEOS2) molecules density by tessellation-based automatic segmentation of super resolution images to determine the cluster size under mock, -Fe supplemented with Ferrozine (-FeFZ) at 100 µM or flg22 at 1µM for 20 minutes. **b**, Quantification of the cluster size in *pUBQ10::SRF3$^{WT}$-mEOS2* (SRF3$^{WT}$-mEOS2) and *pUBQ10::SRF3$^{5KR}$-mEOS2* (SRF3$^{5KR}$-mEOS2) under mock, -Fe supplemented with Ferrozine (-FeFZ) at 100 µM or flg22 at 1µM for 20 minutes. Note that the WT data are used in Fig. Sup 4b. [two-way Kruskal-Wallis coupled with post hoc Steel-Dwass-Critchlow-Fligner procedure was performed ($p<0.05$)]. **c**, Live PALM analysis of *pUBQ10::SRF3$^{WT}$-mEOS2* (SRF3$^{WT}$-mEOS2) and *pUBQ10::SRF3$^{5KR}$-mEOS2* (SRF3$^{5KR}$-mEOS2) molecules density by tessellation-based automatic segmentation of super resolution images to determine the cluster size under mock treatment. **d**, Live PALM analysis of *pUBQ10::SRF3$^{5KR}$-mEOS2* (SRF3$^{5KR}$-mEOS2) molecules density by tessellation-based automatic segmentation of super resolution images to determine the cluster size under mock treatment, -Fe supplemented with Ferrozine (-FeFZ) at 100 µM or flg22 at 1µM for 20 minutes.

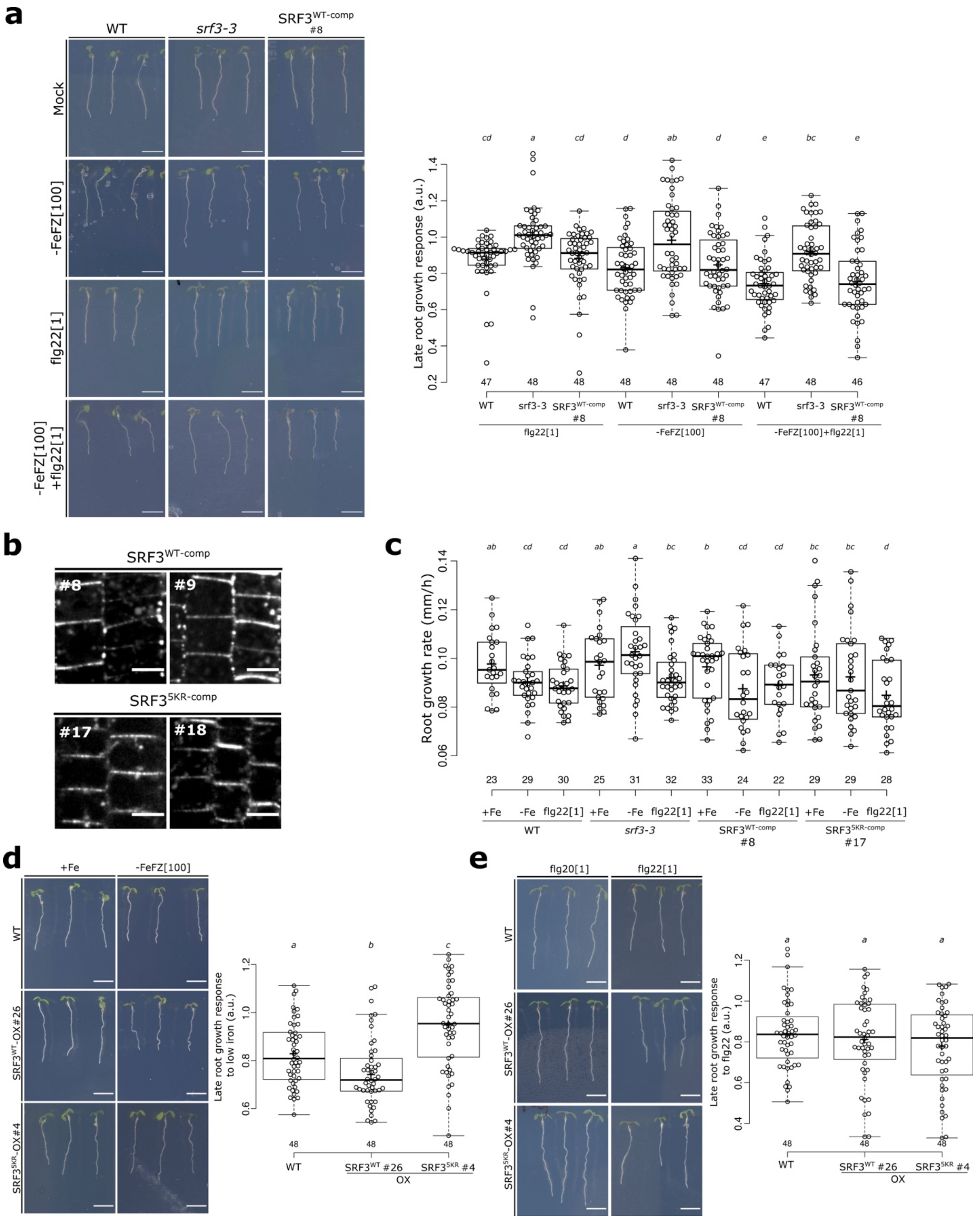

a
WT
srf3-3
SRF3WT-comp #8
Mock
-FeFZ[100]
flg22[1]
-FeFZ[100] +flg22[1]
Late root growth response (a.u.)
cd a cd d ab d e bc e
47 48 48 48 48 48 47 48 46
WT srf3-3 SRF3WT-comp #8 WT srf3-3 SRF3WT-comp #8 WT srf3-3 SRF3WT-comp #8
flg22[1]
-FeFZ[100]
-FeFZ[100]+flg22[1]
b
SRF3WT-comp
#8
#9
SRF35KR-comp
#17
#18
c
Root growth rate (mm/h)
ab cd cd ab a bc b cd cd bc bc d
23 29 30 25 31 32 33 24 22 29 29 28
+Fe -Fe flg22[1] +Fe -Fe flg22[1] +Fe -Fe flg22[1] +Fe -Fe flg22[1]
WT
srf3-3
SRF3WT-comp #8
SRF35KR-comp #17
d
+Fe
-FeFZ[100]
WT
SRF3WT-OX#26
SRF35KR-OX#4
Late root growth response to low iron (a.u.)
a b c
48 48 48
WT SRF3WT #26 SRF35KR #4
OX
e
flg20[1]
flg22[1]
WT
SRF3WT-OX#26
SRF35KR-OX#4
Late root growth response to flg22 (a.u.)
a a a
48 48 48
WT SRF3WT #26 SRF35KR #4
OX

**Figure S7: SRF3 ubiquitination is required to regulate root growth under low iron. a**, Left panel: representative images of 8-day old seedlings of WT, *srf3-3*, *pSRF3::SRF3$^{WT}$-mCITRINE* (*SRF3$^{WT-comp}$*) grown under standard conditions for 5 days and then transferred for 3 days to standard media (mock) or to low iron medium supplemented with Ferrozine at 100µM (–FeFZ[100]) or flg22 at 1 µM or co treated with low iron medium supplemented with Ferrozine at 100µM (–FeFZ[100]) and flg22 at 1 µM. Right panel: quantification of the late root growth response. Numbers indicate independent insertion lines. Scale bar, 1cm. [two-way ANOVA followed by a post hoc Fischer LSD test, letters indicate statistical differences ($p < 0.05$)]. **b**, Confocal images of root epidermal cells of 5-day old seedlings expressing *pSRF3::SRF3$^{WT}$-mCITRINE/srf3-3* (*SRF3$^{WT-comp}$*) and *pSRF3::SRF3$^{5KR}$-mCITRINE/srf3-3* (*SRF3$^{5KR-comp}$*). Numbers indicate independent insertion lines. **c**, Quantification of the root growth rate of WT, *srf3-3*, *pSRF3::SRF3$^{WT}$-mCITRINE/srf3-3* (*SRF3$^{WT-comp}$*) and *pSRF3::SRF3$^{5KR}$-mCITRINE/srf3-3* (*SRF3$^{5KR-comp}$*) under standard media (mock) or low iron (–Fe) or flg22 at 1 µM for 12 hours. Numbers indicate independent insertion lines. [two-way ANOVA followed by a post hoc Fischer LSD T-test, letters indicate statistical differences ($p<0.1$)]. **d**, Left panel: representative images of 8-day old seedlings of WT, *pUBQ10::SRF3$^{WT}$-mCITRINE* (*SRF3$^{WT}$-OX*) or *pUBQ10::SRF3$^{5KR}$-mCITRINE* (*SRF3$^{5KR}$-OX*) grown under standard conditions for 5 days and then transferred for 3 days to standard media (mock) or to low iron medium supplemented with Ferrozine at 100µM (–FeFZ[100]). Right panel: quantification of the late root growth rate (right). Numbers indicate independent insertion lines. Scale bar, 1cm. [two-way ANOVA followed by a post hoc Tukey HSD test, letters indicate statistical differences ($p < 0.05$)]. **e**, Left panel: representative images of 8-day old seedlings of WT, *pUBQ10::SRF3$^{WT}$-mCITRINE* (*SRF3$^{WT}$-OX*) or *pUBQ10::SRF3$^{5KR}$-mCITRINE* (*SRF3$^{5KR}$-OX*) grown under standard conditions for 5 days and then transferred for 3 days to media supplemented with flg20 or flg22 at 1 µM. Right panel: quantification of the late root growth rate (right). Numbers indicate independent insertion lines. Scale bar, 1cm. [two-way ANOVA followed by a post hoc Tukey HSD test, letters indicate statistical differences ($p < 0.05$)].

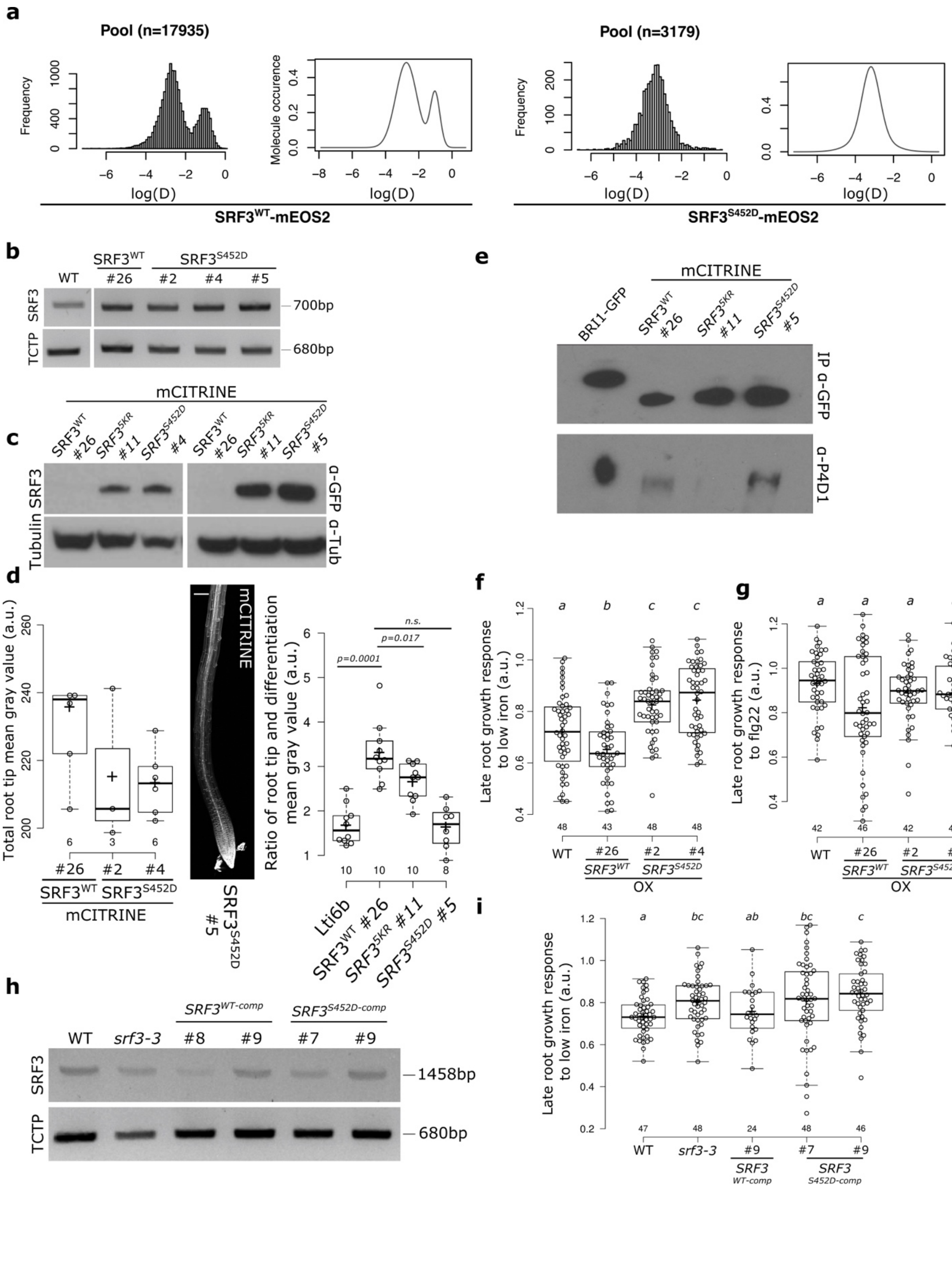
a
Pool (n=17935)
Frequency
Molecule occurence
log(D)
SRF3WT-mEOS2
Pool (n=3179)
SRF3S452D-mEOS2
b
WT
SRF3WT
SRF3S452D
#26
#2
#4
#5
SRF3
TCTP
700bp
680bp
c
mCITRINE
SRF3WT #26
SRF35KR #11
SRF3S452D #4
SRF3S452D #5
SRF3
Tubulin
α-GFP
α-Tub
d
Total root tip mean gray value (a.u.)
mCITRINE
SRF3S452D #5
Ratio of root tip and differentiation mean gray value (a.u.)
p=0.0001
p=0.017
n.s.
Lti6b
SRF3WT #26
SRF35KR #11
SRF3S452D #5
e
mCITRINE
BRI1-GFP
SRF3WT #26
SRF35KR #11
SRF3S452D #5
IP α-GFP
α-P4D1
f
Late root growth response to low iron (a.u.)
WT
SRF3WT
SRF3S452D
OX
g
Late root growth response to flg22 (a.u.)
WT
SRF3WT
SRF3S452D
OX
h
WT
srf3-3
SRF3WT-comp
SRF3S452D-comp
#8
#9
#7
#9
SRF3
TCTP
1458bp
680bp
i
Late root growth response to low iron (a.u.)
WT
srf3-3
SRF3 WT-comp
SRF3 S452D-comp

**Figure S8: Phosphomimic version of flg22-triggered SRF3 phosphorylation prevents its degradation but is still ubiquitinated. a,** Distribution of *pUBQ10::SRF3$^{WT}$-mEOS2* (SRF3$^{WT}$-mEOS2, left) and *pUBQ10::SRF3$^{S452D}$-mEOS2* (SRF3$^{S452D}$-mEOS2, right) molecules according to their apparent diffusion coefficient D obtained by analyzing sptPALM trajectories under mock conditions. Smoothed curves represent the distribution of *pUBQ10::SRF3$^{WT}$-mEOS2* (SRF3$^{WT}$-mEOS2) and *pUBQ10::SRF3$^{S452D}$-mEOS2* (SRF3$^{S452D}$-mEOS2) molecules according to their apparent diffusion coefficient (log(D)) obtained by analyzing the frequency plot using the R mClust package. **b**, RT-PCR using SRF3 primers in *pUBQ10::SRF3$^{WT}$-mCITRINE* (*SRF3$^{WT}$*) and *pUBQ10::SRF3$^{S452D}$-mCITRINE* (*SRF3$^{S452D}$*) of 5-day old seedlings. Numbers indicate independent insertion lines. **c**, Western blot on root material of *pUBQ10::SRF3$^{WT}$-mCITRINE* (*SRF3$^{WT}$*) and *pUBQ10::SRF3$^{5KR}$-mCITRINE* (*SRF3$^{S452D}$*) of 5-day old seedlings using α-GFP and α-TUB. **d**, Left panel: quantification of the mean gray value in the root epidermal cells of 7-day old seedlings expressing *pUBQ10::SRF3$^{WT}$-mCITRINE* (SRF3$^{WT}$) or *pUBQ10::SRF3$^{S452D}$-mCITRINE* (SRF3$^{S452D}$). Numbers indicate independent insertion lines. Middle, confocal image of 7-day old seedlings of roots expressing, *pUBQ10::SRF3$^{S452D}$-mCITRINE* (*SRF3$^{S452D}$*) and the related quantification of the ratio of the root tip and differentiation zone intensity (a.u.). Note that the data for Lti6b, SRF3$^{WT}$ and SRF3$^{5KR}$ are the same as presented in Fig. S3d as images were acquired in the same experiments. Scale bar, 50µm. [Independent two-way Student's T-test ($p < 0.05$)]. **f-g**, Quantification of the late root growth response to low iron (f) and flg22 (g) in WT, *pUBQ10::SRF3$^{WT}$-mCITRINE* (*SRF3$^{WT}$*) or *pUBQ10::SRF3$^{S452D}$-mCITRINE* (*SRF3$^{S452D}$*). Numbers indicate independent insertion lines. [two-way ANOVA followed by a post hoc Tukey HSD test, letters indicate statistical differences ($p < 0.05$) (f) and [two-way Kruskal-Wallis coupled with post hoc Steel-Dwass-Critchlow-Fligner procedure was performed ($p<0.05$) (g)]. **h**, RT-PCR using SRF3 primers in different insertion lines of *pUBQ10::SRF3$^{WT}$-mCITRINE* (*SRF3$^{WT}$*) or *pUBQ10::SRF3$^{S452D}$-mCITRINE* (*SRF3$^{S452D}$*). of 5-day old seedlings. Numbers indicate independent insertion lines. **i**, Quantification of the late root growth response to low iron in WT*, srf3-3*, *pSRF3::SRF3$^{WT}$-mCITRINE/srf3-3* (*SRF3$^{WT\text{-}comp}$*) and *pSRF3::SRF3$^{S452D}$-mCITRINE/srf3-3* (*SRF3$^{S452D\text{-}comp}$*). Numbers indicate independent insertion lines. [two-way ANOVA followed by a post hoc Fischer LSD test, letters indicate statistical difference ($p=0.05$].

**Supplementary Table 1.** RNAseq data
